# The arrow-of-time in neuroimaging time series identifies causal triggers of brain function

**DOI:** 10.1101/2022.05.11.491521

**Authors:** Thomas A. W. Bolton, Dimitri Van De Ville, Enrico Amico, Maria G. Preti, Raphaël Liégeois

**Affiliations:** Connectomics Laboratory, Department of Radiology, Centre Hospitalier Universitaire Vaudois, Switzerland; Department of Clinical Neurosciences, Centre Hospitalier Universitaire Vaudois, Switzerland; Institute of Bioengineering, Center for Neuroprosthetics, École Polytechnique Fédérale de Lausanne, Switzerland; Department of Radiology and Medical Informatics, University of Geneva, Switzerland; CIBM Center for Biomedical Imaging, Switzerland

**Keywords:** Causality, brain function, arrow-of-time, brain dynamics

## Abstract

Moving from *association* to *causal* analysis of neuroimaging data is crucial to advance our understanding of brain function. The arrow-of-time (AoT), *i.e*., the known asymmetric nature of the passage of time, is the bedrock of causal structures shaping physical phenomena. However, almost all current time series metrics do not exploit this asymmetry, probably due to the difficulty to account for it in modelling frameworks. Here, we introduce an AoT-sensitive metric that captures the intensity of causal effects in multivariate time series, and apply it to high-resolution functional neuroimaging data. We find that that causal effects underlying brain function are more clearly localized in space and time than functional activity or connectivity, thereby allowing us to trace neural pathways recruited in different conditions. Overall, we provide a mapping of the causal brain that challenges the association paradigm of brain function.

## Introduction

The advent of functional neuroimaging has provided us with unique insight into the complex spatiotemporal structure of brain function^1^. This organization is classically characterized on the basis of association assessments such as functional connectivity^2^ that was shown to reflect, *e.g*., cognitive status^3,4^ and disease^5–7^. However, the usefulness of this approach has been increasingly questioned as it bears crucial limits in understanding neural communication and pathways^8,9^. Therefore, it is crucial to move from association to causal frameworks to improve the interpretation of functional neuroimaging datasets^10^.

Various approaches have been proposed to extract causal structure from functional imaging time series. They include dynamic causal modelling^11,12^, multivariate autoregressive modelling^13,14^, Granger causality^15,16^, and more application-oriented variants of these^17^. Most, however, do not directly exploit the known asymmetric nature of the passage of time, also called the *arrow-of-time*^18^ (AoT, Fig. 1A). Since the *cause and effect* pattern fundamentally builds upon the AoT, we hypothesize that defining AoT-sensitive metrics of neuroimaging time series will provide unique insights into the causal structure of brain function.

**Figure 1:**
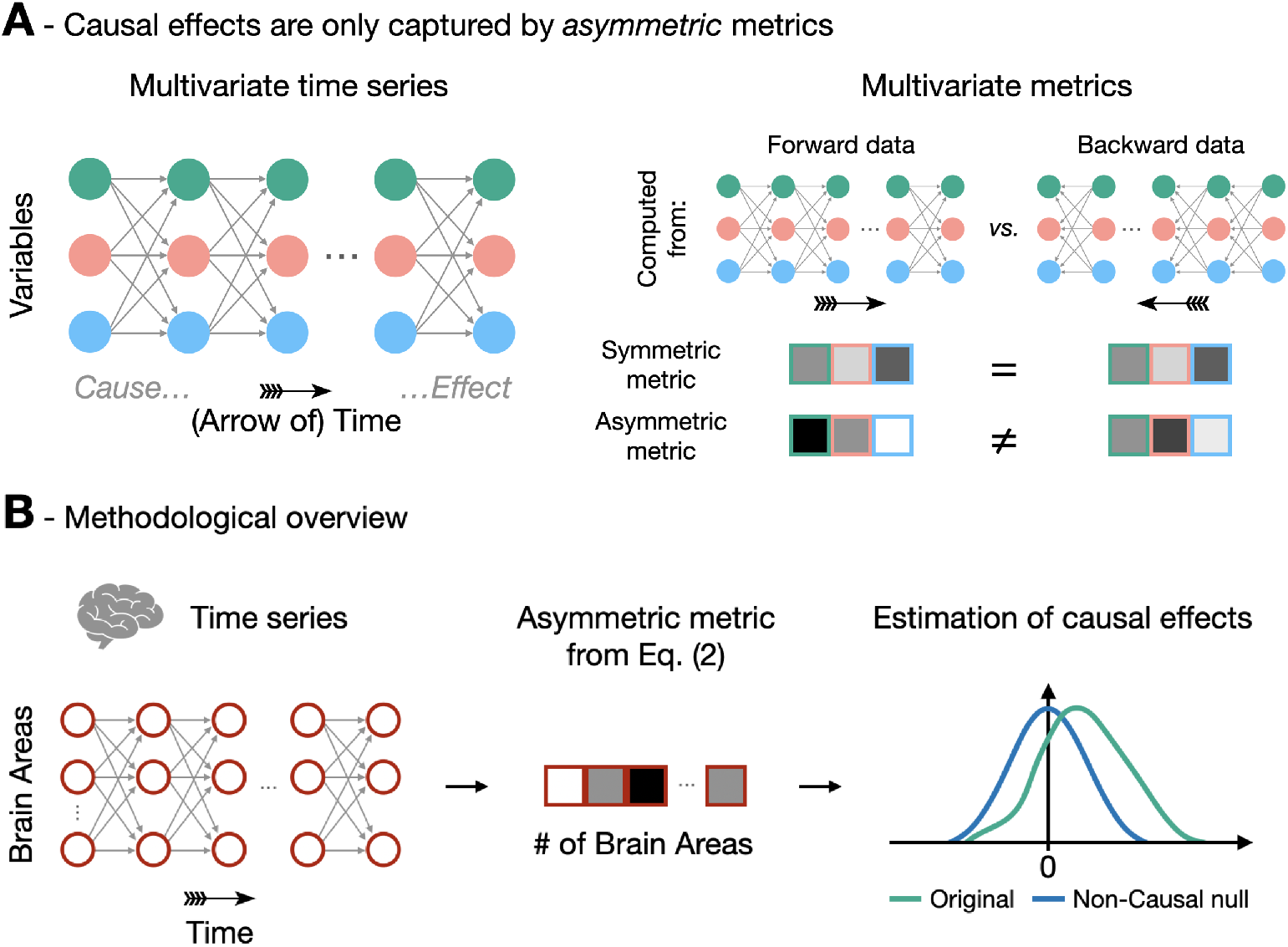
Identifying causal effects in neuroimaging time series using the arrow-of-time. *A*- Since cause precedes effect, causal effects in multivariate time series cannot be identified from metrics that are blind to the AoT. Such symmetric metrics, *e.g*., mean or average correlation over time points, are equal in forward and backward data. In contrast, asymmetric metrics are different in forward and backward data as they are sensitive to the arrow-of-time, thereby bearing the potential of capturing causal effects. *B* - We use fMRI time series acquired during resting state and seven different tasks. The AoT signature is evaluated in these time series using Eq. (2), and the amplitude of the causal effect is assessed by comparison against null time series with no causal effects.

To test this, we introduce a new AoT-sensitive multivariate metric and apply it to high-resolution functional magnetic resonance imaging (fMRI) time series from the Human Connectome Project^19^ (HCP). This metric is a multivariate extension of a previously defined measure^20^, and relies on the comparison of residuals of linear models identified from forward vs backward time series. More precisely, we define *τ*, the AoT strength, as the difference between non-Gaussianity of the residuals of multivariate autoregressive models of forward time series and backward time series (Fig. 1B & Eq. (2), details in *Materials and Methods*). These residuals are expected to be less Gaussian when computed from forward time series^21^, hence we expect *τ* to be positive. This metric is applied on fMRI data from 100 subjects in the resting state and when performing seven different tasks, thereby providing the AoT strength in each brain region, each condition, and as a function of time during paradigms.

We find that in almost all conditions, the AoT strength averaged over brain regions is positive, i.e., the AoT is detected in fMRI time series and shapes their dynamics. Then, we show that patterns of brain regions acting as causal triggers or targets are more localized in space and time as compared to classical activity or connectivity patterns, complementing the ‘networked-brain’ paradigm that has emerged in recent years^22^. Finally, the temporal fluctuations of *τ* during a task paradigm allowed us to identify a causal pathway of neural activations supporting the task. Overall, our results provide unique insight into the causal structure of brain function by leveraging the asymmetric nature of the passage of time to which almost all classical functional neuroimaging metrics are blind^23^.

## Results

### The AoT characterizes cognitive status

We first evaluate *τ* in all conditions as a function of the number of time points used. The AoT strength was computed for each brain region across 100 folds in which subjects were randomly ordered and their time courses were concatenated. The median across folds was taken as an estimate of regional AoT strength, and averaging was then performed across regions to derive a whole-brain AoT heuristic, referred to as 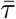. Fig. 2 (top) 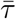 shows as a function of the total amount of considered samples and for all paradigms. In the resting state case (left panel), 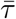 progressively increased as more time points were included, and started to plateau from *n_s_* = 8000 samples, at 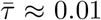. Thus, when sufficient data is available, the AoT is detected in resting-state fMRI time series, confirming the presence of an underlying causal structure.

**Figure 2:**
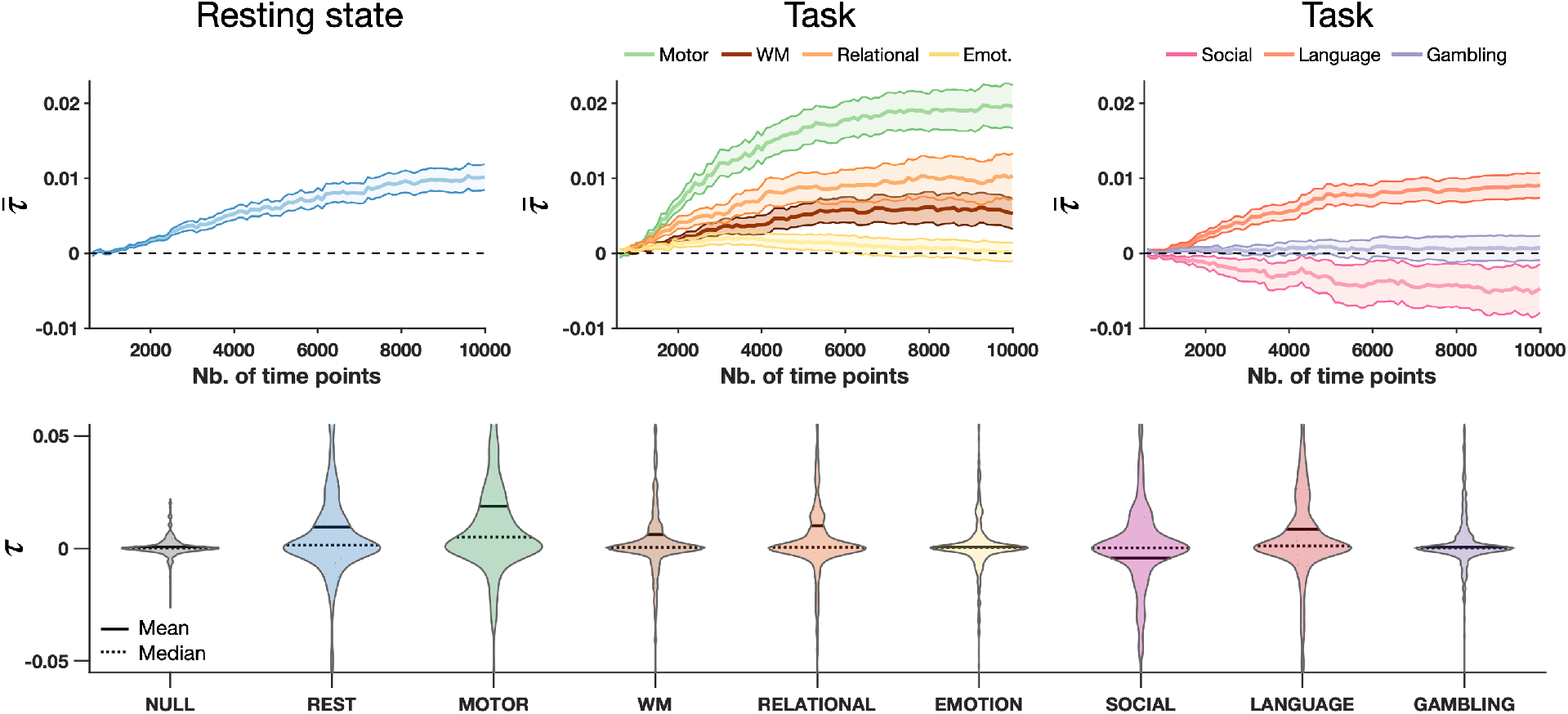
The arrow-of-time is detected in functional magnetic resonance imaging time series. *Top* - Estimated AoT strength across regions 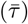 as a function of the number of available samples at rest (left) and for seven different tasks (center, right), with central lines denoting the mean over regions of interest, and surfaces the standard error of the mean. *Bottom* - Distribution of *τ* across regions using *n_s_* = 8000 time points for estimation in non-causal surrogate data (shown here, for indicative purposes, when derived from resting state time courses), at rest, and in seven tasks. Emot.: emotion. WM: working memory.

For task paradigms (middle and right panels), 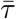 also progressively stabilized as more samples were used, but the asymptotic values differed from case to case: while no sizeable 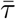 was detected for the gambling (purple) and emotion (yellow) tasks, it was negative for the social task (pink), and positive for the others at varying intensities. The largest AoT was obtained for the motor task, at 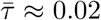. Thus, whole-brain AoT strength also varies as a function of the cognitive task being performed. The negative AoT found in the social task is surprising and suggests that a model assumption has been violated, *e.g*., the presence of an important non-observed variable, or spatial variation in hemodynamic delays.

For subsequent analyses, we focused on the results obtained using 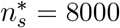 samples, as AoT convergence is observed with this amount of data. Fig. 2 (bottom) shows estimated AoT strength *τ* across regions as a violin plot for each paradigm, as well as when quantified from surrogate data having underwent amplitude-adjusted phase randomization^24^, *i.e*., non-causal null data. In the null case, *τ* was close to zero for all regions, spanning a narrower range of values than for any paradigm. With the exception of the emotion and gambling tasks, while median *τ* across regions was close to zero, mean *τ* was not, denoting that the aforementioned whole-brain causal effects are induced by a subset of brain areas.

### Mapping the causal brain

To determine which brain regions exhibit a significant AoT, we compared them to their respective non-causal null distributions^24^. Fig. 3A shows the results at rest (left), and for the motor task when analyzing full recordings (center) or only task epochs (i.e., having excluded baseline periods, right). Fig. 3B summarizes network contributions to causal effects in all paradigms where contributions to positive and negative *τ* were distinguished. From Eq. (2), it is observed that a positive *τ* corresponds to the presence of a causal *sink, i.e*., the variable is the target of the causal effect. By symmetry, we associate negative values of *τ* to the presence of a causal *source, i.e*., the variable triggers the causal effect (details on the interpretation of positive and negative AoT values are found in the *Materials and Methods*).

**Figure 3:**
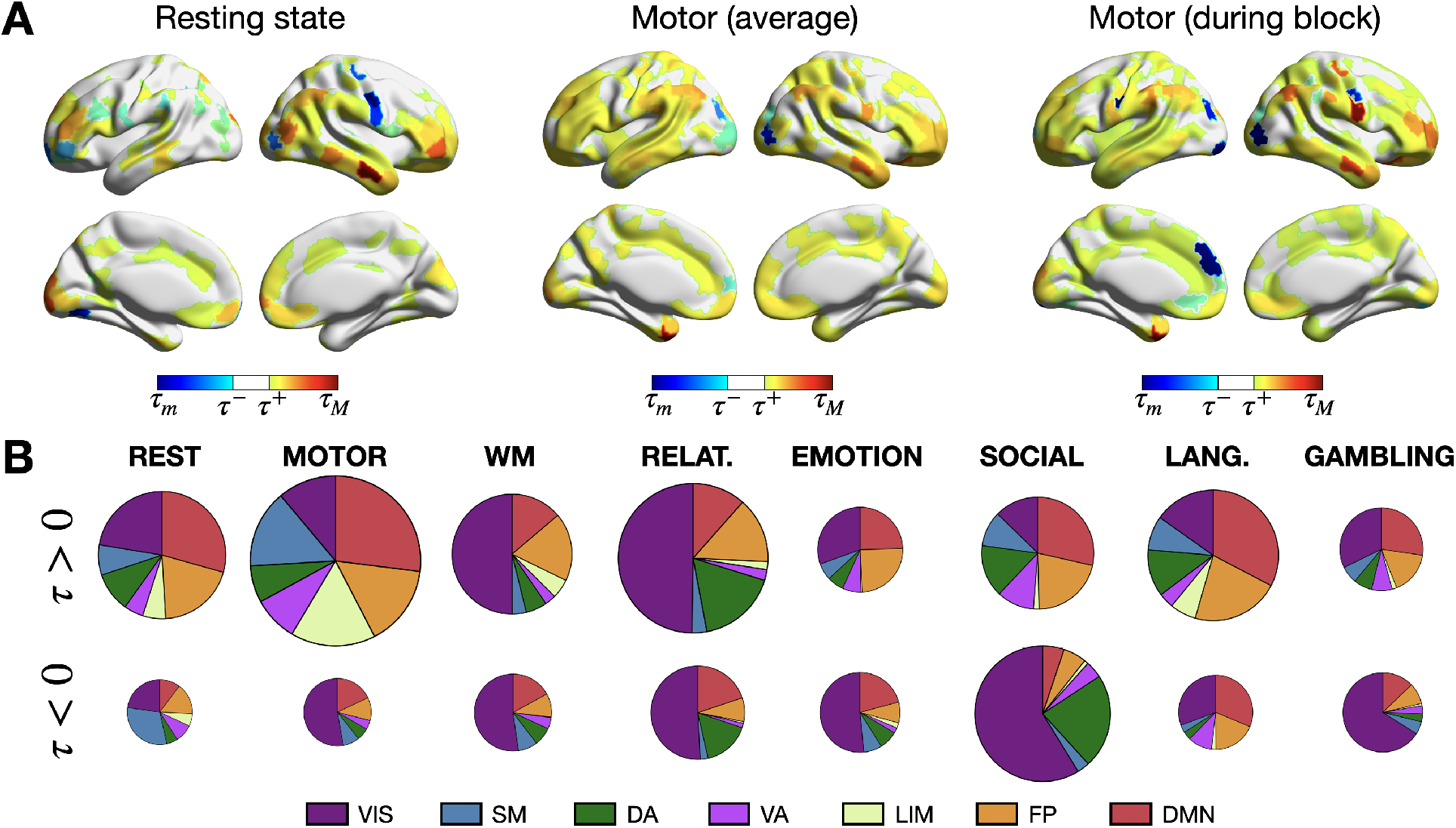
Distinct regional arrow-of-time patterns are observed across paradigms. A - At rest (left), for the full motor task (middle) and when only motor task epochs are considered (right), significant regions in terms of AoT strength. *τ_m_* (*τ_M_*): minimum (maximum) value of *τ, τ*^-^ *τ*^+^): lower (upper) significance threshold at *p* = 0.05 using Bonferroni correction. *B* - For each analyzed paradigm, respective contribution of each of seven canonical networks^25^, shown separately for positive-valued and negative-valued *τ*. All areas (including non-significant ones) are included in this representation. The size of a pie chart is proportional to overall AoT strength in the paradigm at hand.

At rest, 184 regions (43.91%) showed a significant AoT, with a mild right lateralization, and positive-valued *τ* dominated (130 to 54 negative values). The most influential areas primarily spanned the temporal, prefrontal and parietal cortices, and belonged to the default mode and fronto-parietal control networks. Some canonical hubs of these high-level networks showed little significance, such as the posterior cingulate cortex. During the motor task, 284 regions (67.78%) displayed significant causal effects, with no lateralization, and positive values still dominated (214 to 70 negative values). Contributions from the limbic and somatomotor networks were seen in addition to the default mode and fronto-parietal control ones. When excluding baseline moments, 333 regions (79.47%) became significant, with no evident lateralization, and positive values continued to be more prominent (237 to 96 negative ones). Contributions within the somatomotor cortical stripe became stronger, and some other areas with marked negative values were also newly resolved with regard to the two above cases, such as a low-level visual region (R218, *VIS18*) and a prefrontal region (R178, *PFC13*). Overall, these result support the presence of stronger causal mechanisms when a subject engages into the motor task as compared to the resting state.

More broadly across all task paradigms (Fig. 3B), negative-valued *τ* was consistently primarily observed within the visual network, indicating that it always acts as a causal trigger (note that this effect is not observed at rest). This network was also dominant in terms of positive contributions for the working memory and the relational tasks, indicating that it also acts as a causal target in these tasks.

### From causal maps to neural mechanisms

The differences found between full and task-only recordings (Fig. 3A, middle-right) hint at strong temporal fluctuations of the AoT. To ascertain this, we performed a sliding window analysis on the motor task paradigm with a window width of *W* = 20 time points slid by one sample until a full AoT strength time course is computed for each region, and using data from all 100 subjects (Fig. 4A, top). Obtained results were contrasted to the activity time courses temporally smoothed with a moving average filter of length *W*, and to dynamic functional connectivity time courses generated using identical window settings and Pearson’s correlation coefficient as functional connectivity measure. In this latter case, we derived a regional measure by summing all functional connections of an area to the rest of the brain within each temporal window.

**Figure 4:**
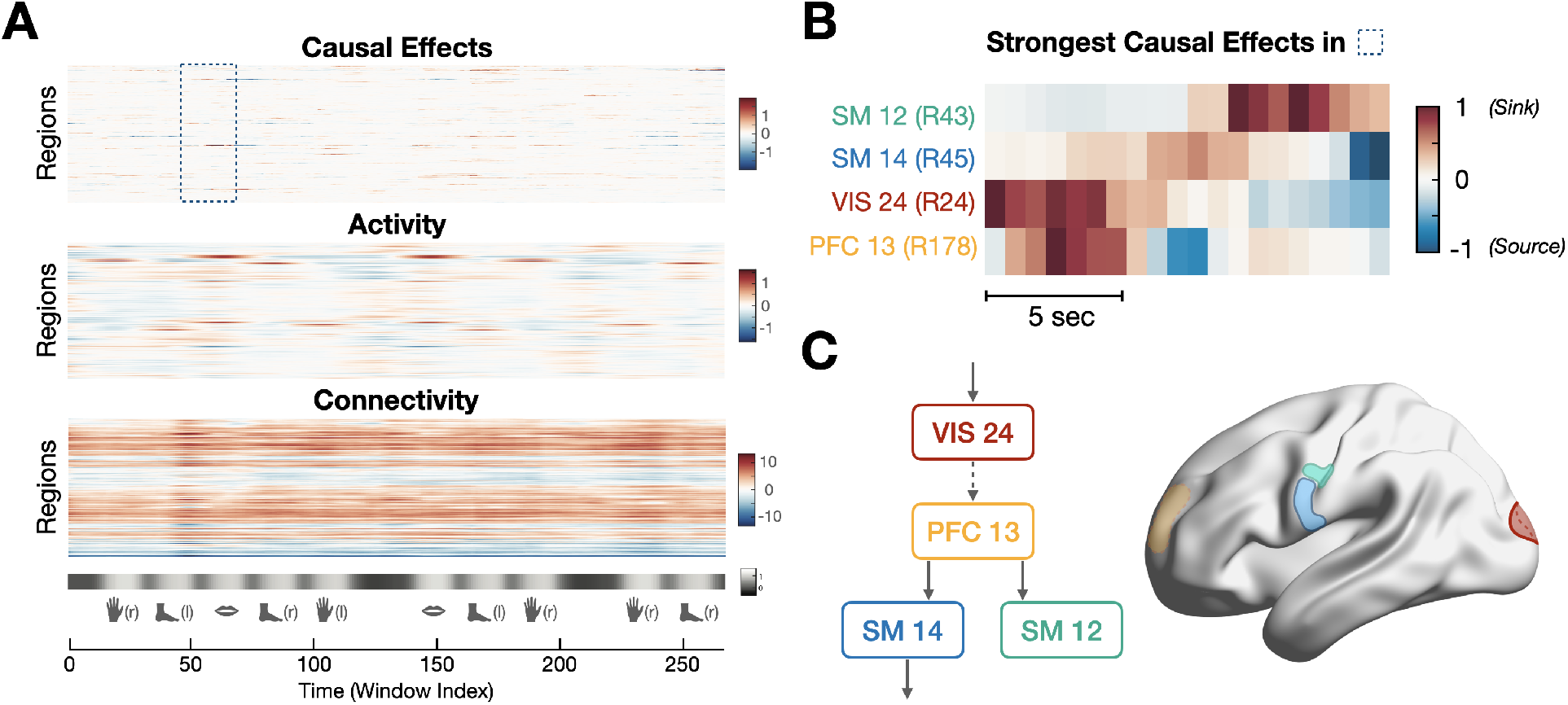
The arrow-of-time identifies spatiotemporally localized causal effects in the motor task. *A* - Measures of causal effects (*τ*, top), activity (middle), and connectivity (bottom) during the motor task paradigm. The paradigm consists of movement epochs (left and right hands and feet, tongue), separated by resting blocks. *B* - Detailed view of causal effects in left hemispheric brain regions showing the strongest causal effects in the interval highlighted in panel A (tongue movement). Positive values suggest that the region acts as a sink for causal effects, while negative values suggest that the region acts as a source of causal effects. *C* - Visualization of the four brain regions in panel B, together with a putative causal pathway recruited when the subjects start moving their tongue.

As expected, clear increases in activity occurred during each of the task epochs in motor regions subserving hand, foot or tongue movement. Connectivity of a given region to the rest of the brain was consistently either positive (denoting a temporally stable regime with more prominent correlation to the rest of the brain), or negative (more prominent anti-correlation). On the whole, activity and connectivity fluctuations were relatively diffuse in time (spanning full task epochs) and in space (involving many different areas). In contrast, causal effect time courses were highly localized in space (typically only applying to individual regions at any given time point), and occurred within shorter time intervals with fast transition from positive (causal target) to negative (causal source) values.

Fig. 4B exemplifies the evolution of causal effects when transiting from baseline to the first tongue movement epoch (see highlighted area in panel A, bottom), for the four left hemispheric brain regions with the largest extent of temporal fluctuations of *τ* within this interval. Consistent with the paradigm’s demands, these regions were motor (*SM12* and *SM14*, for tongue movement), visual (*VIS24*, for parsing the provided instructions), and prefrontal (*PFC13*, to trigger movement execution). When the visual cue is provided to the subjects, *VIS24* becomes a causal sink. Shortly afterwards, *PFC13* becomes a sink, as visual information is treated frontally to make the decision to move. This information is then transmitted to the rest of the brain, as *PFC13* becomes a causal source (see the temporally localized negative values in its time course), while *SM14* and, later on, *SM12* become sinks. Finally, *SM14* further transmits the information and becomes a source to trigger motion. Fig. 4C schematically summarizes these observations.

## Discussion

Here, we introduced a new AoT-sensitive metric that captures causal effects in multivariate time series. Applied to fMRI data, we showed that causal effects (i) shape brain function in all conditions, (ii) are highly localized in space and time, and (iii) reflect underlying neural mechanisms. These results are found to be robust to head motion, to the use of a different metric of non-Gaussianity, and to varying processing strategies (see *Supplementary Material*). While causality has been assessed in neuroscience and neuroimaging using other methods^17,26–28^, this is to the best of our knowledge the first use of the AoT to interrogate causality in neuroimaging data, thereby providing a new and natural description of the causal brain.

### The AoT provides a new perspective into the causal structure of time series

The term ‘arrow-of-time’ has been coined by Sir A. Eddington almost a century ago to *express this one-way property of time which has no analogue in space*^18^. Rather surprisingly, identifying the AoT from time series is not trivial and most current AoT detection methods rely on deep learning^29–31^. Other approaches instead exploit simpler features such as the distribution^20^ or the independence^32^ of linear model residuals in forward and backward time series. The latter measures, from which we defined *τ* in Eq. (2), also come with a natural interpretation in terms of causality as they leverage causal inference theory to detect the AoT^21,32^. Therefore, the interpretation of *τ* in terms of causality comes with all causal inference assumptions and guarantees, which is not necessarily the case of other causality detection methods used in neuroimaging studies that encode different forms of causality^23,33^.

Identifying causal effects rather than association effects in multivariate time series comes with estimation challenges. For example, it is seen from Fig. 2 (see also *Supplementary Material* for further evidence) that at least ~ 1000 fMRI time points are required to identify stable AoT patterns. In contrast, stable patterns of functional connectivity, *i.e*., of correlation, can be identified from as little as around 100 fMRI time points^34^. Exploiting the non-Gaussianity of time series through kurtosis also requires cautious estimation of group effects as this metric relates to outliers in a distribution. For this reason, we took several precautions to maximize the stability of our maps: we evaluated our group (original and null) results from the *median* over folds (thus accounting for the selection of different subjects and making our results more generalizable), and adopted the most efficient sample selection scheme after evaluating several candidates (see *Supplementary Material*). Resorting to non-Gaussianity of linear models was important in order to unambiguously identify causal structures; indeed, linear-Gaussian approaches usually only lead to a *class* of possible models, a.k.a. Markov equivalence class, equivalent in their conditional correlation structure and from which no unique causal structure can be inferred^21,35^.

### The association brain vs the causal brain

The current perception of brain function has been built from association metrics of functional neuroimaging data, thus probing the ‘association brain’. For example, functional connectivity^2,36,37^ canonical resting-state networks^1,25^, and most representations of brain dynamics such as (innovation-driven) co-activation patterns^38,39^, dynamic modes^40^, or sliding window-based states^41–43^ are defined from association metrics, *e.g*., correlation, which are blind to causality. By leveraging advances in causal inference, we defined a simple metric that exploits time series asymmetry induced by causal effects. This shift of the methodological paradigm lays the ground to a shift of canonical representations of brain function and dynamics. Furthermore, a causal representation of brain function also comes with promises for the cognitive and clinical use of neuroimaging data as the causal brain is expected to reflect underlying neural mechanisms^9^, as illustrated in Fig. 4B/C. Recent neuroimaging endeavours further substantiate this potential: after training a deep learning network to distinguish between temporal segments of forward and backward fMRI time series, Deco et al.^31^ not only observed a variable AoT strength (inferred from classification accuracy on unseen data) across cognitive states, but also between healthy subjects and patients suffering from bipolar disorder, attention deficit hyperactivity disorder or schizophrenia. In another study leveraging the same framework on electrocorticography data, de la Fuente et al.^44^ also revealed that deep sleep and ketamine-induced anesthesia lowered the differences between forward time series and their inverted counterparts, i.e., decreased AoT strength.

Our results show that the topology of the the causal brain exhibits strong differences as compared to the association brain. Specifically, the dynamic tracking of the AoT in Fig. 4A revealed how remarkably localized it was with regard to functional activation and connectivity. While these two common measures reflect the overall simultaneity in activation across regions, when information has already arrived and been locally amplified (for instance, somatomotor areas in our motor task example), our AoT metric captures the arrival and departure of information. It thus more finely pinpoints the spatial entry and exit points of neural pathways, as well as their exact temporality. As a consequence, time-averaged representations of the causal brain might be harder to interpret as they destroy the rich temporal structure of causal effects (Fig. 3A). In particular, further work will be required to efficiently characterize the causal brain, *e.g*., through causal networks accounting for its specificities. Finally, the present association vs causal brain dichotomy differs from the one between functional and effective connectivity^2^. Indeed, using the current causal inference nomenclature, most functional and effective connectivity measures would be classified as association measures (see, e.g., the discussion on the nature of Granger causality in Pearl et al.^23^, Chap. I).

### Limitations and further considerations

The proposed characterization of causal effects comes with the assumptions and limitations of the modelling framework in Eqs. (1)-(2). In particular, we limit our assessment to linear and non-Gaussian causal effects. This is motivated by the indeterminacy inherent to linear-Gaussian assessments^21^, but does not mean that causal effects cannot be Gaussian. Future work will explore whether relaxing these assumptions, e.g., using convergent cross mapping^45^ or other nonlinear approaches^46^, provides new insights into the causal brain. Robustness to violation of causal sufficiency, i.e., the presence of non-observed variables, would also need to be further assessed^47,48^, potentially by including additional experimental variables of interest such as a record of the visual cue or electrophysiological variables. Then, comparisons across paradigms must be interpreted with caution as while the total number of samples was the same, the length of the paradigms was different. Thus, a distinct number of subjects contributed to the estimates in each case. This directly relates to the question of individual as opposed to population-wise causal effects, and further work will explore the potential of the causal brain as a subject-level marker^49,50^. Finally, our framework is directly applicable to other neuroimaging modalities, e.g., electro- or magneto-encephalography, but also outside of neuroimaging to any multivariate time series dataset.

### Conclusion

Together, our findings suggest that a causal assessment of neuroimaging data indeed provides new insights into the neural mechanisms underlying brain function. More precisely, our mapping of the causal brain hints at key differences as compared to association paradigms of brain function during rest and task, *e.g*., in terms of spatial and temporal localization. In light of this, brain imaging studies have an opportunity to move beyond classical association paradigms and unveil information contained in neuroimaging data to which current metrics are blind.

## Materials and Methods

### Data acquisition and preprocessing

We considered *S* = 100 unrelated healthy subjects from the Human Connectome Project S900 data release (46 males, 54 females, mean age = 29.1 ± 3.7 years). We used fMRI recordings acquired at rest and during 7 tasks (emotion, gambling, language, motor, relational, social, working memory), for which ethical approval was obtained within the HCP. Our analyses focused on the first of two available resting state sessions, and on each available task session, purely on the left-right phase encoding direction runs. Right-left phase encoding data were examined in supplementary analyses (see *Supplementary Material*).

To generate regional fMRI time courses, for each run of interest, minimally preprocessed data from the HCP^19,51^ were taken as input. Nuisance signals were first removed from the voxel-wise fMRI time courses, including linear and quadratic trends, the six motion parameters and their first derivatives, as well as the average white matter and cerebrospinal fluid signals and their first derivatives. In our main analyses, the global signal was also included as a confounding variable. In additional analyses (see *Supplementary Material*), we contrasted the obtained results to those without global signal regression, and also examined the impacts of performing scrubbing as a final preprocessing step. Voxel-wise time courses were averaged within each region of a parcellation containing 400 cortical^52^ and 19 subcortical^51,53^ areas, for a total of *R* = 419 parcels, and eventually z-scored. To complement these analyses, we also considered cortical atlases containing 200 and 800 regions ^52^ (see *Supplementary Material*).

### AoT quantification

To quantify AoT strength across brain regions, we extend a univariate metric defined previously^20^ to the multivariate case. First, we fit a first-order multivariate autoregressive model to concatenated fMRI time series population-wise^54^, both in the *forward* and in the *backward* directions as shown in Eq. (1):

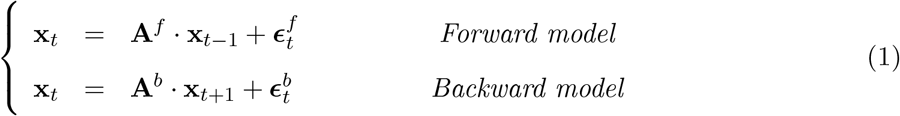

where **x***_t_* is of size *R* × 1, **A***^f^* and **A***^b^* each have size *R* × *R*, and the residuals 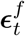 and 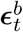 are of size *R* × 1. The model parameters are estimated using ordinary least squares^55^, and successive samples that originate from separate subjects (owing to the concatenation step) are excluded. Then, the presence of causal effects in different brain regions is assessed by comparing non-Gaussianity of forward and backward residuals. This was motivated by the fact that residuals of linear models of true cause-effect links (in this case, the forward model) are more non-Gaussian than the residuals of the reversed linear models (in this case, the backward model)^21^. Concretely, with *T* the total number of time points, we define 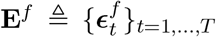 and 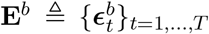 as the forward and backward error distributions. Regional AoT strength *τ*(*i*) is then estimated as:

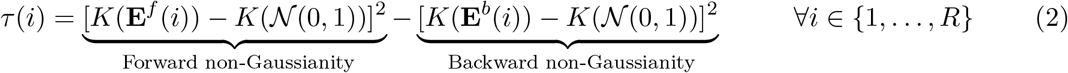

where *K*(·) denotes the *kurtosis* of a distribution, and 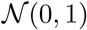 stands for the standard normal distribution. In the case of a marked AoT, non-Gaussianity of residuals is larger in the forward than in the backward model, and *τ*(*i*) is positive. Region *i* is then a causal *sink*, primarily receiving information from the rest of the brain. By symmetry, we say that if *τ*(*i*) is negative, brain region i is a causal *source*. Note, however, that a negative value of *τ* suggests that one model assumption has been violated, *e.g*., due to the presence of an unobserved variable, or due to different delays in hemodynamic responses, and interpretation of negative values of *τ*(*i*) should be cautious. Finally, we also devised an alternative metric relying on the Kullback-Leibler divergence to quantify AoT strength (see *Supplementary Material* for details).

### Regional AoT patterns

Using 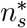 samples, regional AoT patterns were extracted for each paradigm of interest. For the compatible tasks, the same process was also conducted after the removal of baseline epochs. To do so, individual binarized paradigm time courses (0=rest, 1=task) were convolved with the canonical haemodynamic response function from SPM12, and resulting time points with a value larger/lower than 0.5 were treated as task/rest samples. Of note, since less samples are then available per subject, the obtained AoT estimates gather data from a more extended set of subjects compared to the full recording case.

To study the contribution of separate networks to the AoT patterns, each cortical brain region was assigned to one of seven canonical whole-brain resting state networks^25^ through a majority voting procedure. Positive- and negative-valued AoT contributions were separately quantified.

### Significance assessment

To assess AoT significance, comparison was performed to null data for which causal effects were destroyed. For this purpose, for each paradigm at hand, amplitude-adjusted phase randomization^24^ was applied to the original time courses to generate *n_n_* = 100 null realizations. We considered this surrogate procedure in order to destroy causal effects while preserving the original auto-correlation structure and sampling distribution, including potential non-Gaussian effects. For each set of null data, using 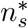 samples, AoT strength was calculated across 100 folds, and the median was taken as an estimate of null regional AoT strength. The mean and standard deviation were quantified for each regional null distribution, and *τ* was deemed significant if it exceeded the Bonferroni-corrected 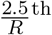 or 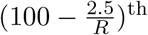 null percentiles (*τ*^-^ and *τ*^+^ in Fig. 3, respectively).

### Software availability

All the scripts used in this work were implemented and tested in MATLAB, versions 2014b, 2020b and 2021a (MathWorks, Natick, MA, USA). They can be freely downloaded from the following link: https://github.com/TiBiUan/AoT_Benchmarking.git. For figure generation, we used the cbrewer and *BrainNet Viewer*^56^ (version 1.7) utilities.

## Acknowledgements

RL acknowledges support by the Swiss National Centre of Competence in Research - Evolving Language (grant number 51NF40_180888). MGP was supported by the CIBM Center for Biomedical Imaging, a Swiss research center of excellence founded and supported by Lausanne University Hospital (CHUV), University of Lausanne (UNIL), Ecole Polytechnique Fédérale de Lausanne (EPFL), University of Geneva (UNIGE) and Geneva University Hospitals (HUG).

## Notes

### Competing Interest Statement

The authors have declared no competing interest.

## References

[1] Damoiseaux, J., Rombouts, S., Barkhof, F., Scheltens, P., Stam, C., Smith, S.M., et al. Consistent restingstate networks across healthy subjects. Proceedings of the national academy of sciences 2006;103(37):13848–13853.

[2] Friston, K.J.. Functional and effective connectivity: a review. Brain connectivity 2011;1(1):13–36.

[3] Greicius, M.D., Krasnow, B., Reiss, A.L., Menon, V.. Functional connectivity in the resting brain: a network analysis of the default mode hypothesis. Proceedings of the National Academy of Sciences 2003;100(1):253–258.

[4] van den Heuvel, M.P., Mandl, R.C.W., Kahn, R.S., Hulshoff Pol, H.E.. Functionally linked resting-state networks reflect the underlying structural connectivity architecture of the human brain. Human brain mapping 2009;30(10):3127–41. doi:10.1002/hbm.20737.

[5] Drysdale, A.T., Grosenick, L., Downar, J., Dunlop, K., Mansouri, F., Meng, Y., et al. Resting-state connectivity biomarkers define neurophysiological subtypes of depression. Nat Med 2017;23(1):28–38. doi:10.1038/nm.4246.

[6] Bassett, D.S., Nelson, B.G., Mueller, B.A., Camchong, J., Lim, K.O.. Altered resting state complexity in schizophrenia. Neuroimage 2012;59(3):2196–207. doi:10.1016/j.neuroimage.2011.10.002.

[7] Anderson, J.S., Nielsen, J.A., Froehlich, A.L., DuBray, M.B., Druzgal, T.J., Cariello, A.N., et al. Functional connectivity magnetic resonance imaging classification of autism. Brain 2011;134(Pt 12):3742–54. doi:10.1093/brain/awr263.

[8] Reid, A.T., Headley, D.B., Mill, R.D., Sanchez-Romero, R., Uddin, L.Q., Marinazzo, D., et al. Advancing functional connectivity research from association to causation. Nat Neurosci 2019;22(11):1751–1760. doi:10.1038/s41593-019-0510-4.

[9] Weichwald, S., Peters, J.. Causality in cognitive neuroscience: Concepts, challenges, and distributional robustness. Journal of Cognitive Neuroscience 2021;33(2):226–247. doi:https://doi.org/10.1162/jocn_a_01623.

[10] Siddiqi, S.H., Kording, K.P., Parvizi, J., Fox, M.D.. Causal mapping of human brain function. Nature Reviews Neuroscience 2022;URL: https://doi.org/10.1038/s41583-022-00583-8. doi:10.1038/s41583-022-00583-8.

[11] Friston, K.J., Harrison, L., Penny, W.. Dynamic causal modelling. Neuroimage 2003;19(4):1273–1302.

[12] Friston, K.. Causal modelling and brain connectivity in functional magnetic resonance imaging. PLoS biol 2009;7(2):e1000033.

[13] Valdés-Sosa, P.A., Sánchez-Bornot, J.M., Lage-Castellanos, A., Vega-Hernández, M., Bosch-Bayard, J., Melie-García, L., et al. Estimating brain functional connectivity with sparse multivariate autoregression. Philos Trans R Soc Lond B Biol Sci 2005;360(1457):969–81. doi:10.1098/rstb.2005.1654.

[14] Rogers, B.P., Katwal, S.B., Morgan, V.L., Asplund, C.L., Gore, J.C.. Functional mri and multivariate autoregressive models. Magnetic resonance imaging 2010;28(8):1058–1065.

[15] Barnett, L., Seth, A.K.. The mvgc multivariate granger causality toolbox: a new approach to granger-causal inference. J Neurosci Methods 2014;223:50–68. doi:10.1016/j.jneumeth.2013.10.018.

[16] Barrett, A.B., Barnett, L., Seth, A.K.. Multivariate granger causality and generalized variance. Phys Rev E Stat Nonlin Soft Matter Phys 2010;81(4 Pt 1):041907. doi:10.1103/PhysRevE.81.041907.

[17] Seth, A.K., Barrett, A.B., Barnett, L.. Granger causality analysis in neuroscience and neuroimaging. J Neurosci 2015;35(8):3293–7. doi:10.1523/JNEUROSCI.4399-14.2015.

[18] Eddington, A.S.. The Nature of the Physical World-Chap. V. Cambridge University Press; 1928.

[19] Van Essen, D.C., Smith, S.M., Barch, D.M., Behrens, T.E., Yacoub, E., Ugurbil, K., et al. The wu-minn human connectome project: an overview. Neuroimage 2013;80:62–79.

[20] Hernández-Lobato, J., Morales-Mombiela, P., Suárez, A.. Gaussianity measures for detecting the direction of causal time series. IJCAI International Joint Conference on Artificial Intelligence 2011;:1318–1323doi:10.5591/978-1-57735-516-8/IJCAI11-223.

[21] Shimizu, S., Hoyer, P.O., Hyvarinen, A., Kerminen, A.. A linear non-gaussian acyclic model for causal discovery. Journal of Machine Learning Research 2006;7(72):2003–2030. URL: http://jmlr.org/papers/v7/shimizu06a.html.

[22] Betzel, R.F., Bassett, D.S.. Multi-scale brain networks. Neuroimage 2017;160:73–83. doi:10.1016/j.neuroimage.2016.11.006.

[23] Pearl, J.. Causality: Models, reasoning, and inference, second edition. Causality 2000;29. doi:10.1017/CBO9780511803161.

[24] Theiler, J., Eubank, S., Longtin, A., Galdrikian, B., Farmer, J.D.. Testing for nonlinearity in time series: the method of surrogate data. Physica D: Nonlinear Phenomena 1992;58(1):77–94.

[25] Yeo, B.T.T., Krienen, F.M., Sepulcre, J., Sabuncu, M.R., Lashkari, D., Hollinshead, M., et al. The organization of the human cerebral cortex estimated by intrinsic functional connectivity. Journal of neurophysiology 2011;106:1125–1165. doi:10.1152/jn.00338.2011.

[26] Friston, K., Moran, R., Seth, A.K.. Analysing connectivity with granger causality and dynamic causal modelling. Curr Opin Neurobiol 2013;23(2):172–8. doi:10.1016/j.conb.2012.11.010.

[27] Roebroeck, A., Formisano, E., Goebel, R.. The identification of interacting networks in the brain using fmri: Model selection, causality and deconvolution. Neuroimage 2011;58(2):296–302. doi:10.1016/j.neuroimage.2009.09.036.

[28] Cekic, S., Grandjean, D., Renaud, O.. Time, frequency, and time-varying granger-causality measures in neuroscience. Stat Med 2018;37(11):1910–1931. doi:10.1002/sim.7621.

[29] Wei, D., Lim, J.J., Zisserman, A., Freeman, W.T.. Learning and using the arrow of time. In: Proceedings of the IEEE Conference on Computer Vision and Pattern Recognition (CVPR). 2018,.

[30] Seif, A., Hafezi, M., Jarzynski, C.. Machine learning the thermodynamic arrow of time. Nature Physics 2020;17(1):105–113. URL: https://doi.org/10.10382Fs41567-020-1018-2. doi:10.1038/s41567-020-1018-2.

[31] Deco, G., Perl, Y., Sitt, J., Tagliazucchi, E., Kringelbach, M.. Deep learning the arrow of time in brain activity: characterising brain-environment behavioural interactions in health and disease. bioRxiv 2021;.

[32] Bauer, S., Schölkopf, B., Peters, J.. The arrow of time in multivariate time series. Proceedings of Machine Learning Research 2016;48:2043–2051.

[33] White, H., Chalak, K., Lu, X.. Linking granger causality and the pearl causal model with settable systems. In: Popescu, F., Guyon, I., editors. Proceedings of the Neural Information Processing Systems Mini-Symposium on Causality in Time Series; vol. 12 of Proceedings of Machine Learning Research. Vancouver, Canada: PMLR; 2011, p. 1–29. URL: https://proceedings.mlr.press/v12/white11.html.

[34] Van Dijk, K.R.A., Hedden, T., Venkataraman, A., Evans, K.C., Lazar, S.W., Buckner, R.L.. Intrinsic functional connectivity as a tool for human connectomics: theory, properties, and optimization. J Neurophysiol 2010;103(1):297–321. doi:10.1152/jn.00783.2009.

[35] Spirtes, P., Glymour, C., Scheines, R.. Causation, Prediction, and Search. The MIT Press; 2000. ISBN 978-1-4612-7650-0. doi:10.1007/978-1-4612-2748-9.

[36] Power, J.D., Cohen, A.L., Nelson, S.M., Wig, G.S., Barnes, K.A., Church, J.A., et al. Functional network organization of the human brain. Neuron 2011;72(4):665–678.

[37] Biswal, B., Yetkin, F.Z., Haughton, V.M., Hyde, J.S.. Functional connectivity in the motor cortex of resting human brain using echo-planar mri. Magnetic resonance in medicine 1995;34:537–541.

[38] Liu, T.T.. Noise contributions to the fmri signal: An overview. Neuroimage 2016;143:141–151. doi:10.1016/j.neuroimage.2016.09.008.

[39] Karahanoğlu, F.I., Van De Ville, D.. Transient brain activity disentangles fmri resting-state dynamics in terms of spatially and temporally overlapping networks. Nat Commun 2015;6:7751. doi:10.1038/ncomms8751.

[40] Casorso, J., Kong, X., Chi, W., Van De Ville, D., Yeo, B.T.T., Liégeois, R.. Dynamic mode decomposition of resting-state and task fmri. Neuroimage 2019;194:42–54. doi:10.1016/j.neuroimage.2019.03.019.

[41] Allen, E.A., Damaraju, E., Plis, S.M., Erhardt, E.B., Eichele, T., Calhoun, V.D.. Tracking whole-brain connectivity dynamics in the resting state. Cerebral cortex 2014;24(3):663–676. doi:10.1093/cercor/bhs352.

[42] Preti, M.G., Bolton, T.A., Van De Ville, D.. The dynamic functional connectome: State-of-the-art and perspectives. Neuroimage 2017;160:41–54. doi:10.1016/j.neuroimage.2016.12.061.

[43] Lurie, D.J., Kessler, D., Bassett, D.S., Betzel, R.F., Breakspear, M., Kheilholz, S., et al. Questions and controversies in the study of time-varying functional connectivity in resting fmri. Network Neuroscience 2020;4(1):30–69.

[44] de la Fuente, L.A., Zamberlan, F., Bocaccio, H., Kringelbach, M.L., Deco, G., Perl, Y.S., et al. Temporal irreversibility of neural dynamics as a signature of consciousness. bioRxiv 2021;.

[45] Sugihara, G., May, R., Ye, H., Hsieh, C.h., Deyle, E., Fogarty, M., et al. Detecting causality in complex ecosystems. Science 2012;338(6106):496–500.

[46] Runge, J., Nowack, P., Kretschmer, M., Flaxman, S., Sejdinovic, D.. Detecting and quantifying causal associations in large nonlinear time series datasets. Science Advances 2019;5(11).

[47] Zhang, J.. On the completeness of orientation rules for causal discovery in the presence of latent confounders and selection bias. Artificial Intelligence 2008;172(16):1873–1896. URL: http://www.sciencedirect.com/science/article/pii/S0004370208001008. doi:https://doi.org/10.1016/j.artint.2008.08.001.

[48] Runge, J.. Causal network reconstruction from time series: From theoretical assumptions to practical estimation. Chaos: An Interdisciplinary Journal of Nonlinear Science 2018;28(7):075310. URL: https://doi.org/10.1063/1.5025050. doi: 10.1063/1.5025050. arXiv:https://doi.org/10.1063/1.5025050.

[49] Finn, E.S., Shen, X., Scheinost, D., Rosenberg, M.D., Huang, J., Chun, M.M., et al. Functional connectome fingerprinting: identifying individuals using patterns of brain connectivity. Nat Neurosci 2015;18(11):1664–1671. URL: http://dx.doi.org/10.1038/nn.4135. doi:10.1038/nn.4135.

[50] Van De Ville, D., Farouj, Y., Preti, M.G., Liégeois, R., Amico, E.. When makes you unique: Temporality of the human brain fingerprint. Sci Adv 2021;7(42):eabj0751. doi:10.1126/sciadv.abj0751.

[51] Glasser, M.F., Sotiropoulos, S.N., Wilson, J.A., Coalson, T.S., Fischl, B., Andersson, J.L., et al. The minimal preprocessing pipelines for the human connectome project. Neuroimage 2013;80:105–124.

[52] Schaefer, A., Kong, R., Gordon, E.M., Laumann, T.O., Zuo, X.N., Holmes, A.J., et al. Local-global parcellation of the human cerebral cortex from intrinsic functional connectivity mri. Cereb Cortex 2018;28(9):3095–3114. doi:10.1093/cercor/bhx179.

[53] Fischl, B., Salat, D.H., Busa, E., Albert, M., Dieterich, M., Haselgrove, C., et al. Whole brain segmentation: automated labeling of neuroanatomical structures in the human brain. Neuron 2002;33(3):341–55. doi:10.1016/s0896-6273(02)00569-x.

[54] Liégeois, R., Li, J., Kong, R., Orban, C., Van De Ville, D., Ge, T., et al. Resting brain dynamics at different timescales capture distinct aspects of human behavior. Nature Communications 2019;10(1):2317. doi:10.1038/s41467-019-10317-7.

[55] Stoica, P., Moses, R.L.. Spectral analysis of signals. Pearson/Prentice Hall Upper Saddle River, NJ; 2005.

[56] Xia, M., Wang, J., He, Y.. Brainnet viewer: a network visualization tool for human brain connectomics. PLoS One 2013;8(7):e68910. doi:10.1371/journal.pone.0068910.

