## Supplementary Material for "The arrow-of-time in neuroimaging time series identifies causal triggers of brain function"

### Supplementary Results

#### *Quality of AoT estimation as a function of sample number*

In the resting state case, each available run contains 1200 time points. AoT strength can thus be quantified using yet many more samples than done in our main results. To verify whether the detected causal effects would be altered in such a setting, we estimated $\tau$ at rest using up to 100000 samples (Fig. 1).

As more samples were considered (*i.e.*, moving downward in the top plot), the obtained AoT pattern was further strengthened, and the involved brain regions remained identical. This was confirmed by a spatial correlation between successively estimated patterns that already exceeded

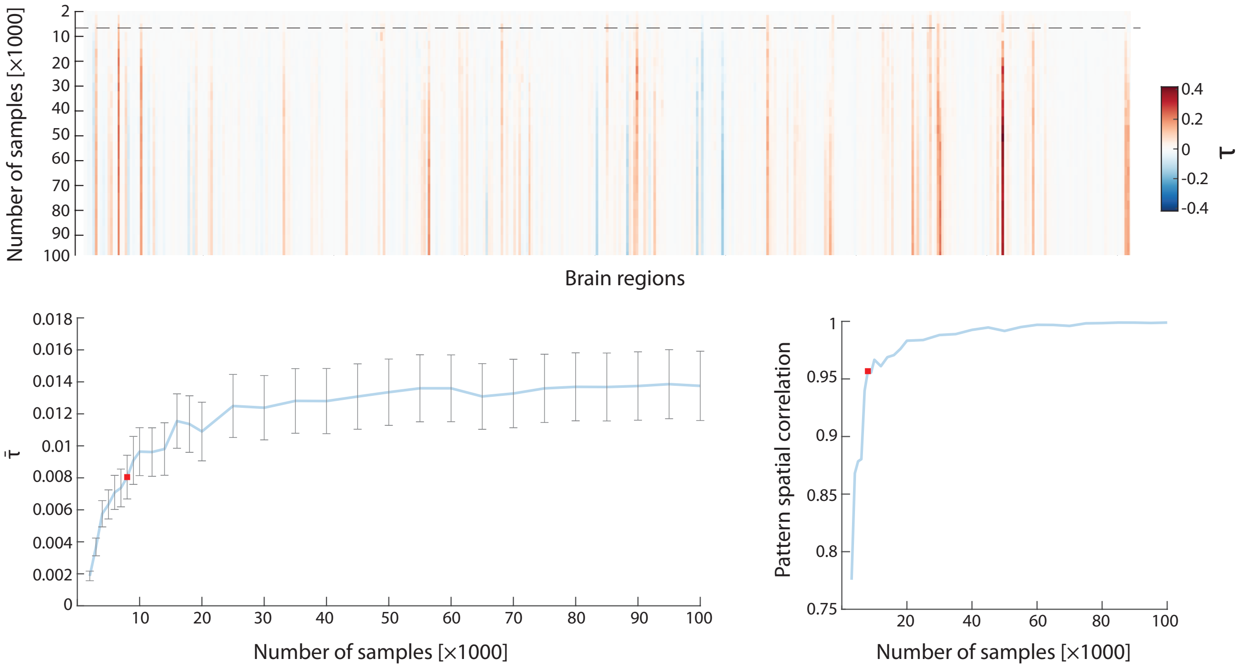


Figure 1: *Top* - Estimation of $\tau$ in the resting state case when from 2000 to 100000 samples are used (top to bottom in the heatmap), for all brain regions (left to right). *Bottom left* - Convergence of mean AoT across regions $\bar{\tau}$ as more samples are considered. Error bars denote standard error of the mean. *Bottom right* - Spatial correlation between the AoT patterns obtained using two successive numbers of samples. The estimates reached using $n_s^* = 8000$ samples (the value selected for main analyses) are highlighted by a dashed horizontal line (top panel) or a red rectangle (bottom panels).

0.95 for $n_s^* = 8000$ samples (bottom right plot). As could be expected given the strengthening of the detected causal effects, $\bar{\tau}$ also continued to moderately increase as further samples were added, until $\bar{\tau} \approx 0.013$ (bottom left plot).

On task paradigms (Fig. 2), similar observations could be made in terms of pattern strengthening and convergence, regardless of the exact task at hand. Thus, using $n_s^* = 8000$ samples to estimate AoT strength appears sufficient, regardless of the investigated paradigm, to detect all the brain regions implicated in causal brain mechanisms.

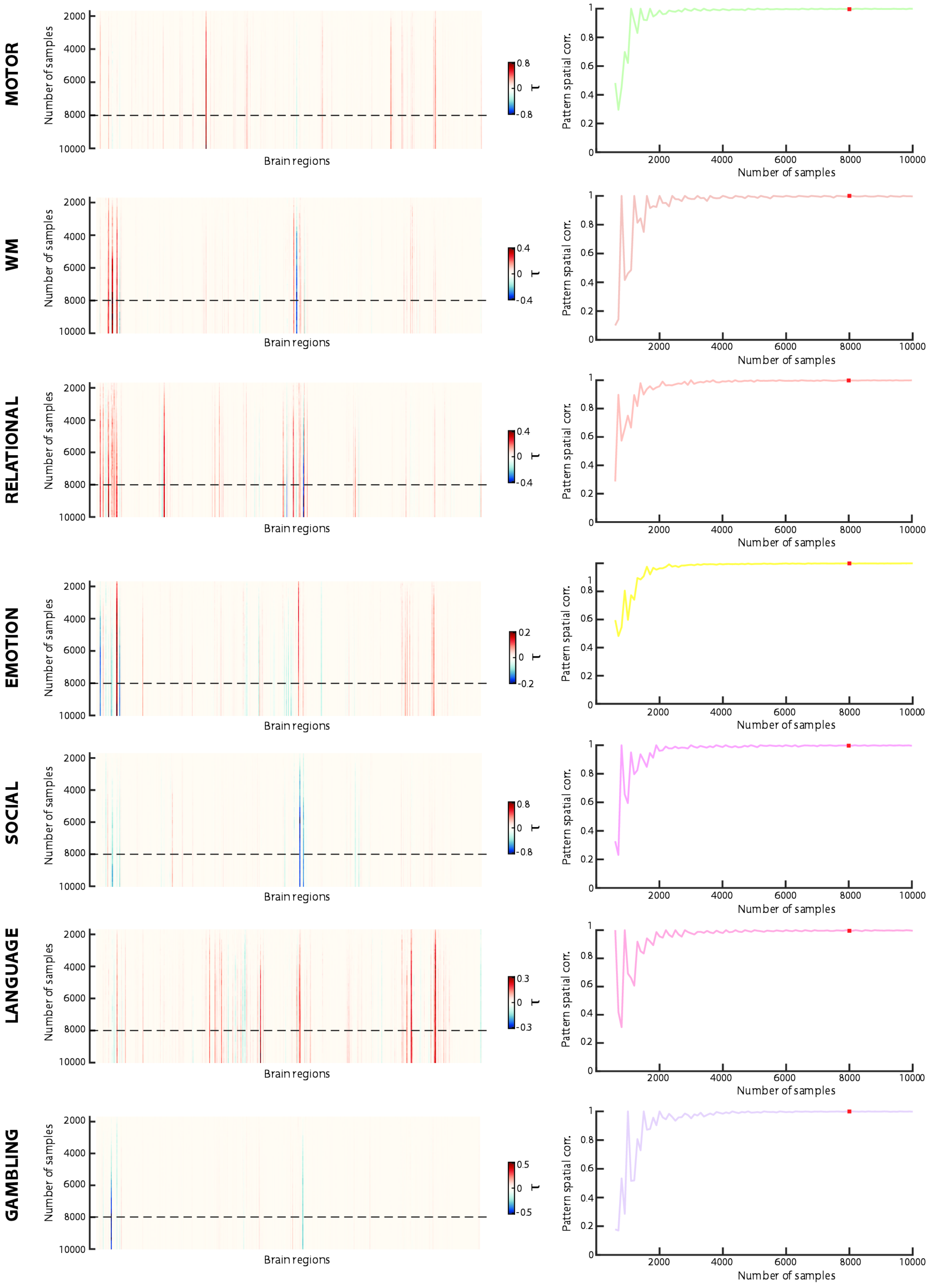


Figure 2: For each task (row of plots), estimation of $\tau$ when up to 10000 samples are used (top to bottom in the heatmaps), for all brain regions (left to right), and spatial correlation between the AoT patterns obtained using two successive numbers of samples. The estimates reached using $n_s^* = 8000$ samples (the value selected for main analyses) are highlighted by a dashed horizontal line (left heatmaps) or a red rectangle (right plots).

*Regional AoT patterns across tasks*

Fig. 3 shows the AoT patterns extracted from full task recordings using $n_s^* = 8000$ samples, when convergence is already achieved as demonstrated above. Below, drawing from past work[1], we first briefly summarize the main components of each task. We then discuss the largest AoT contributors in terms of how they fit each paradigm's demands. Since we consider full paradigms, for which a given area may transit between acting as a causal source or sink over the course of time, we do not take sign into account in what follows.

The working memory task was an $N$-back task in which images of faces, tools, places and body parts were presented to the subjects. Half of the blocks consisted in a 0-back task, and half in a 2-back task.

In terms of AoT strength, the most influential areas were largely confined to the occipital cortex. There were also two anterior frontopolar regions from the right hemisphere (R341, R343), known to be important in working memory tasks for the manipulation of integrated information[2].

In the relational task, for *relational blocks*, the subjects were simultaneously shown two pairs of objects, with each object a combination of a shape and a texture. They had to determine which dimension differed between the top objects, and whether the bottom objects also differed along that same dimension. In *matching blocks*, they were instead shown two objects at the top of the screen, one at the bottom, and a word (either "shape" or "texture") in the middle. They had to determine whether the bottom object matched any of the top ones in terms of the displayed dimension.

Visual regions were the strongest contributors to the AoT pattern. Of all the tasks, this was the one with the broadest array of significant visual contributions. The intraparietal sulcus (IPS) was also resolved bilaterally (R73, R75, R282, R286). The IPS plays a role in multiple object tracking[3] as well as in the short-term memory for multifeature objects[4], both of which are specifically important for this task. The left rostrolateral prefrontal cortex (R135) was detected as well, and contributes to relational integration during reasoning[5].

In the emotion task, in *emotional blocks*, participants were shown one face at the top of the screen, and two at the bottom. They had to determine which of these two matches the top one.

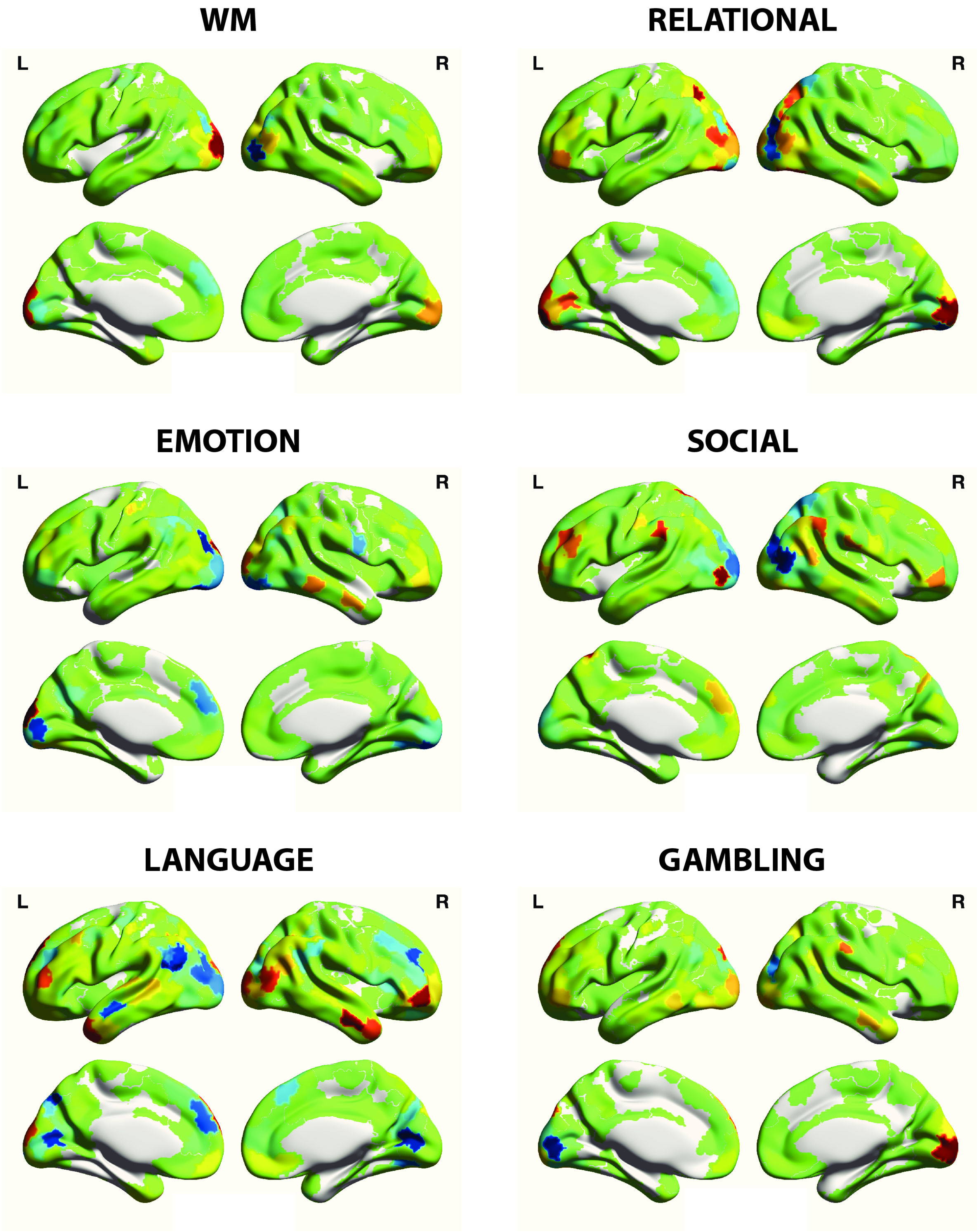


Figure 3: Regional AoT patterns for all tasks (except the motor one, shown in the main results), considering full time courses and using $n_s^* = 8000$ samples for estimation.

The faces had either an angry or a fearful expression. In the *shape blocks*, they instead had to determine which of two bottom shapes matched the top one.

AoT strength was overall low in the emotion task, for which the only few notable regions were part of the visual system.

In the social task, participants were shown movie clips of geometrical shapes that either interacted in a certain way (*social blocks*), or moved randomly (*random blocks*). They had to decide whether the shapes were socially interacting or moving randomly, with the possibility to state that they were unsure.

As in the above cases, the strongest contributors were visual regions. This task was the one with the second broadest set of influential visual areas. Similarly to the relational task, R282 and R286 were detected, which makes sense as the social movie clips also involved multiple objects to track. In addition, an area in the left angular gyrus (R72) previously linked to action awareness representation[6] was pinpointed, as well as the left supramarginal gyrus (R95), which enables to retain an abstract representation of serial order information[7], and the right inferior parietal lobule (R332), implicated in the discrimination of direction changes[8].

In the language task, participants were stimulated auditorily instead of visually. In *story blocks*, they were provided with short stories followed by a 2-alternative forced-choice question about the topic of the story. In the *maths blocks*, they were given a mathematical operation and had to select the correct answer out of two choices.

Fittingly given the auditory nature of stimulation, the language task was the one for which the fewest visual regions were influential (R24, R221 and R222 only). In addition, several areas linked to theory of mind (ToM) were resolved, including the left medial prefrontal cortex (involved in several ToM-related functions[9], R179) and the bilateral temporal pole (directly linked to ToM in story comprehension[10], R124,R367,R368).

In the gambling task, subjects were asked to guess whether the number (between 1 and 9) on a mystery card would be lower or larger than 5. In the *win blocks*, the outcome would be decided so as to favour gains, while in the *loss blocks*, it would instead favour losses. Participants eventually received their total gain as US dollars.

Only a restricted set of visual areas were influential in this task.

In summary, these observations collectively strengthen our main results from Fig. **??** in showing that our AoT-sensitive metric can reveal important brain regions implicated in low-level and high-level brain functions.

#### *Impact of baseline epochs on estimated AoT strength*

When baseline epochs were removed from compatible task paradigms (Figure 4), convergence onto a task-specific AoT pattern was still observed from $\approx$ 2000 samples. For the motor, emotion and social tasks, asymptotic $\bar{\tau}$ values increased in magnitude compared to their respective full run counterparts (compare to Figure **??**), while for the language task, there was no major change, and for the working memory task, there was a switch to negative values.

The pinpointed regional pattern without baseline epochs remained overall similar for the motor and emotion tasks, to the exception of some areas which switched sign (positive to negative $\tau$ in

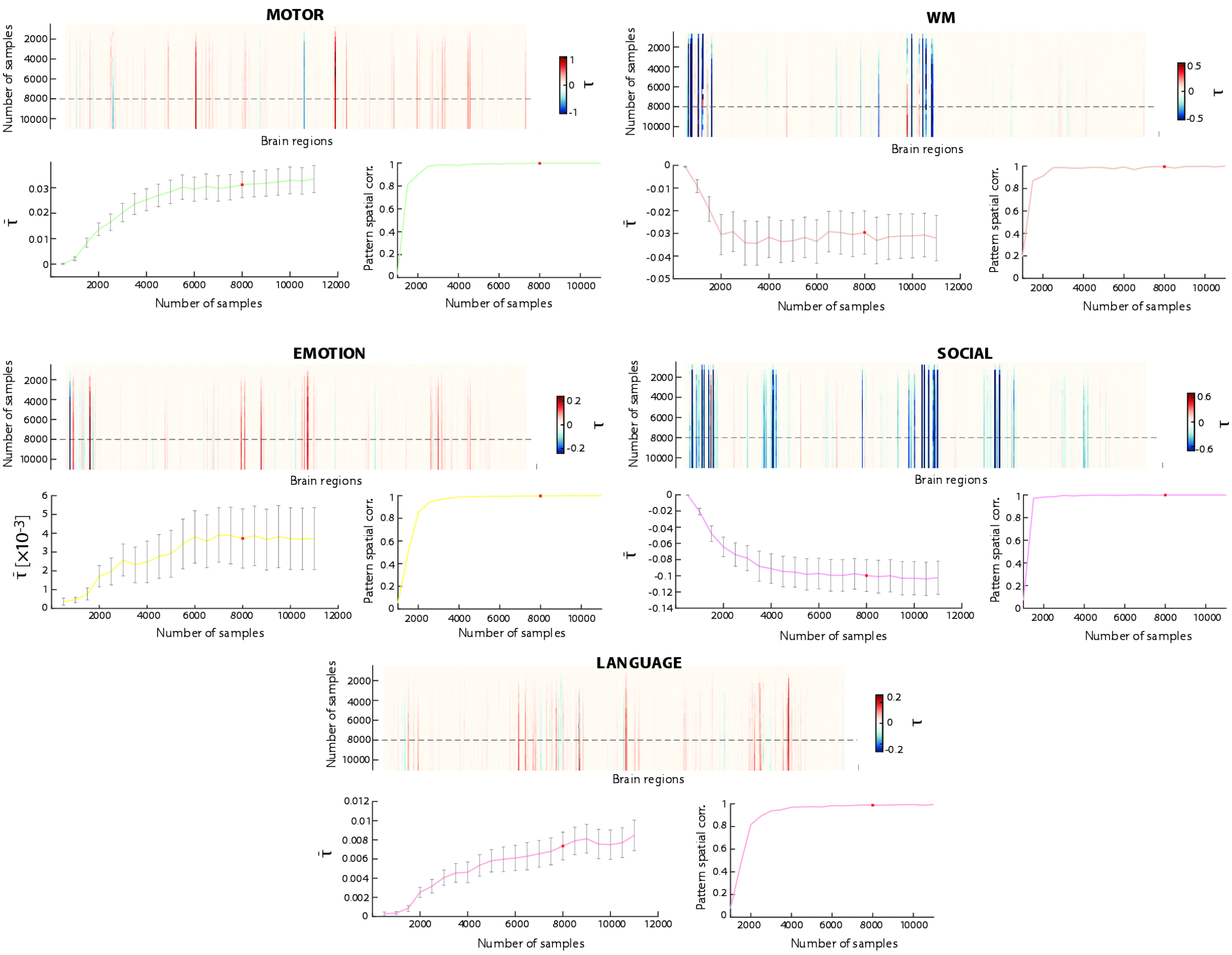


Figure 4: For each task, estimation of $\tau$ when up to 10000 samples are used (top to bottom in the heatmaps), for all brain regions (left to right); convergence of mean AoT across regions $\bar{\tau}$ as more samples are considered (with error bars reflecting standard error of the mean); and spatial correlation between the AoT patterns obtained using two successive numbers of samples. The estimates reached using $n_s^* = 8000$ samples (the value selected for main analyses) are highlighted by a dashed horizontal line (top heatmaps) or a red rectangle (bottom plots).

the former case, and negative to positive $\tau$ in the latter case). This may be because their role as causal source or sink fluctuates as a function of epoch type.

A marked transition to negative-valued $\tau$ was seen in the working memory and social task cases, particularly for visual areas. The working memory task was specifically designed to probe visual function on top of working memory. Similarly, the social task involves particularly salient visual stimulation in the form of moving geometric shapes. Negative-valued $\tau$ when focusing on task epochs highlights that during such a condition, visual regions behave as strong causal sources, transmitting information to the rest of the brain.

For the language task, changes upon removing baseline epochs were minimal. Interestingly, as this is the only task that does not rely on visual stimulation, the changes observed for other tasks are likely largely modulated by the involvement of the visual network.

*Null AoT distributions are similar across paradigms*

In Fig. **??**, the distributions of $\tau$ values across regions were shown across paradigms, and compared to surrogate values derived from one realization of an amplitude-adjusted phase randomization process[11]. For the sake of conciseness, null data were only shown when generated from resting state time courses. Here, we wish to confirm that null distributions are in fact extremely similar regardless of the distorted input data (*i.e.*, resting state or any of the task paradigms).

The distribution of $\tau$ values across regions following amplitude-adjusted phase randomization is shown in Figure 5, for 15 concatenated null realizations, in the resting state case, for the motor task with or without including baseline epochs, and for the other 6 tasks. In all cases, the median (horizontal line) and the mean (empty rectangle) both remained almost equal to zero, and the range of taken values was similar. This confirms that our main results capture significant causal effects in all the investigated paradigms. Note that the range of values is larger here than in Fig. **??** because it only displayed the results for one null realization instead of 15 as here.

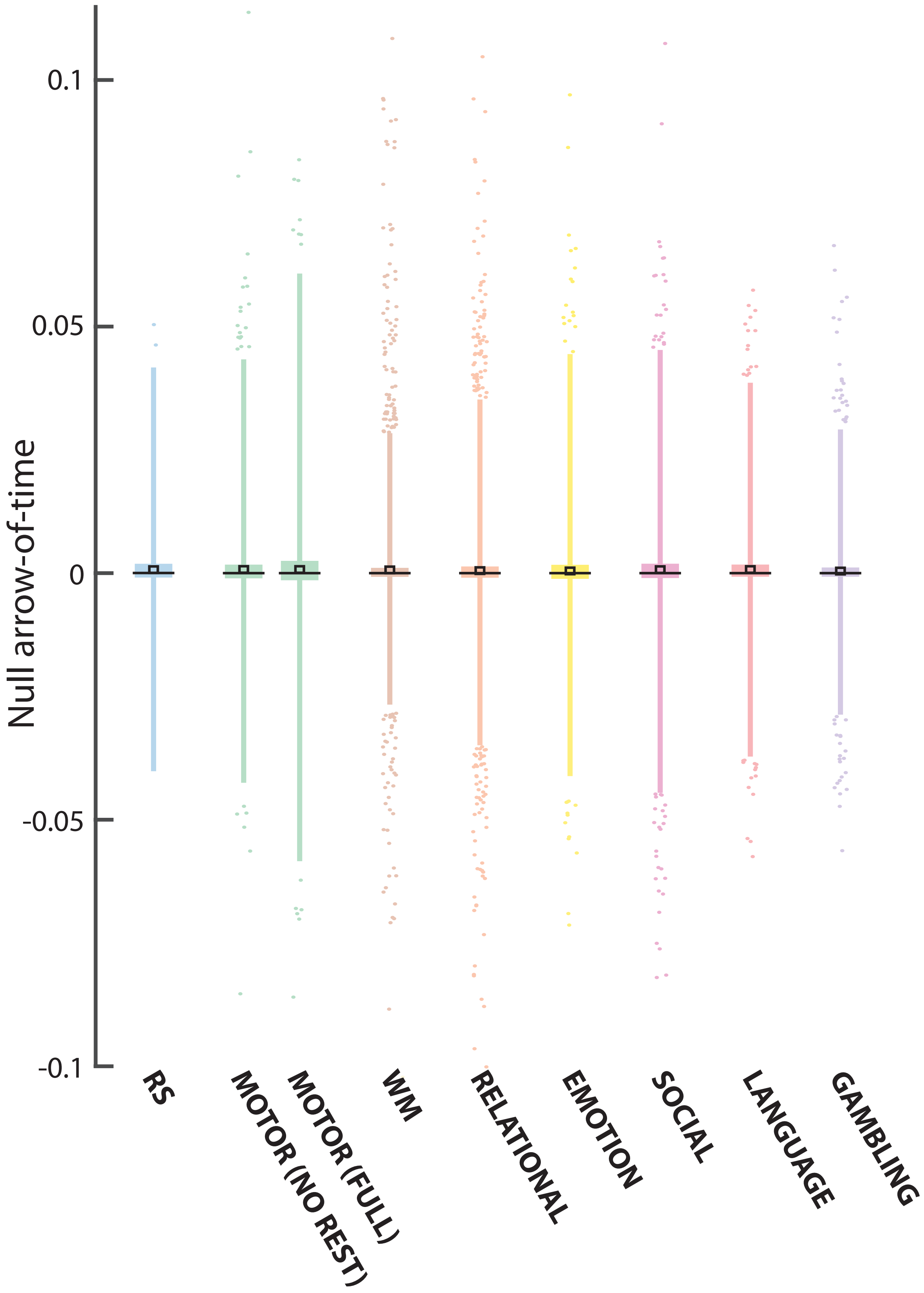


Figure 5: For the resting state (blue), motor (green, with/without baseline epochs on the left/right) and other task paradigms (color coding as in Fig. ??), distribution of null $\tau$ values across regions and 15 null realizations. Note that data points are drawn as outliers if they are larger than $Q_3 + 15 \cdot (Q_3 - Q_1)$ or smaller than $Q_1 - 15 \cdot (Q_3 - Q_1)$, with $Q_1$ and $Q_3$ the 25$^{\text{th}}$ and 75$^{\text{th}}$ percentiles, respectively.

*Dynamic evolution of AoT across tasks*

To complement Fig. ?? in which we focused on the motor task, we provide below similar visualizations of causal effects over time for the other compatible task paradigms.

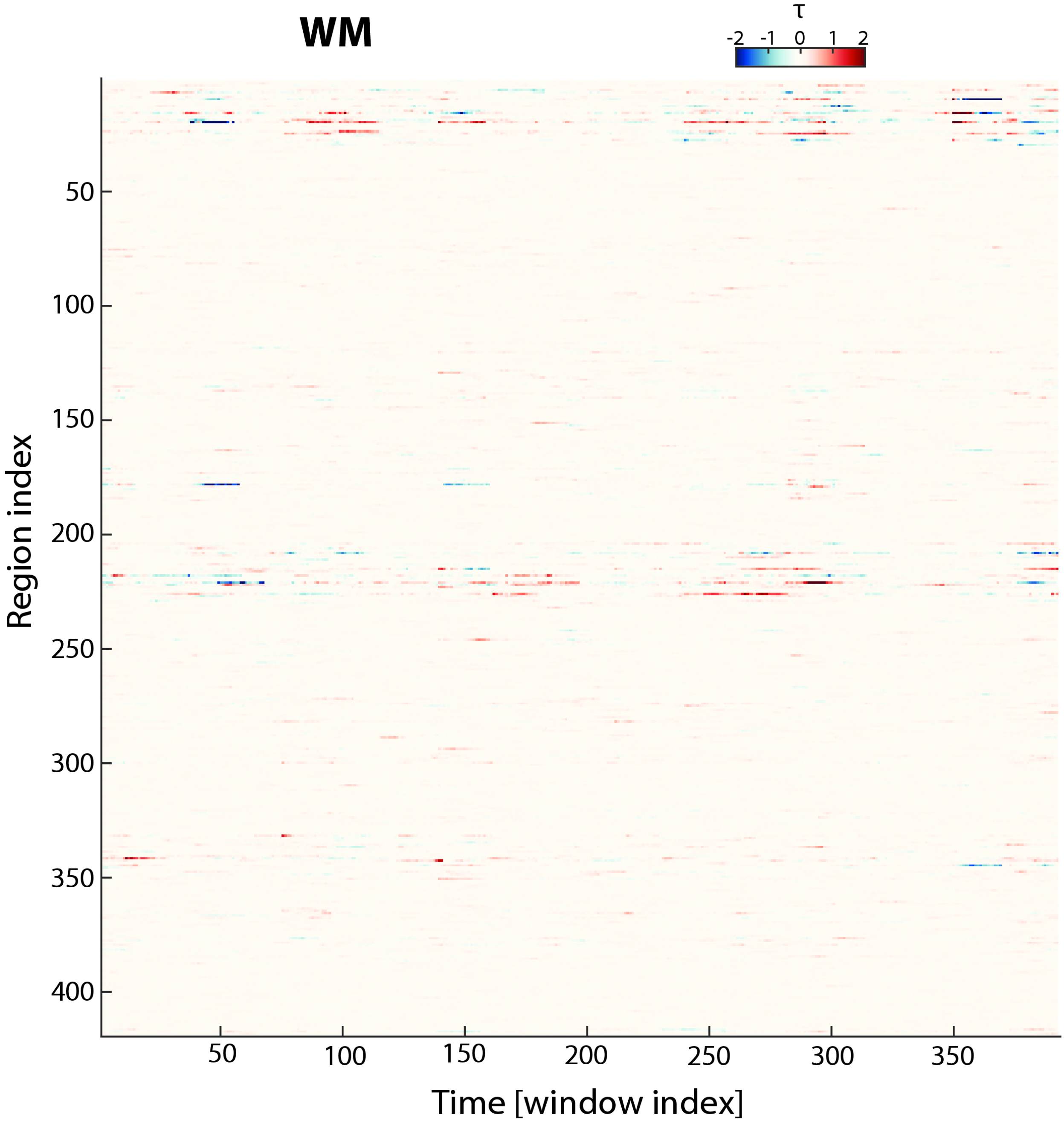


Figure 6: Evolution of causal effects during the working memory task. WM: working memory.

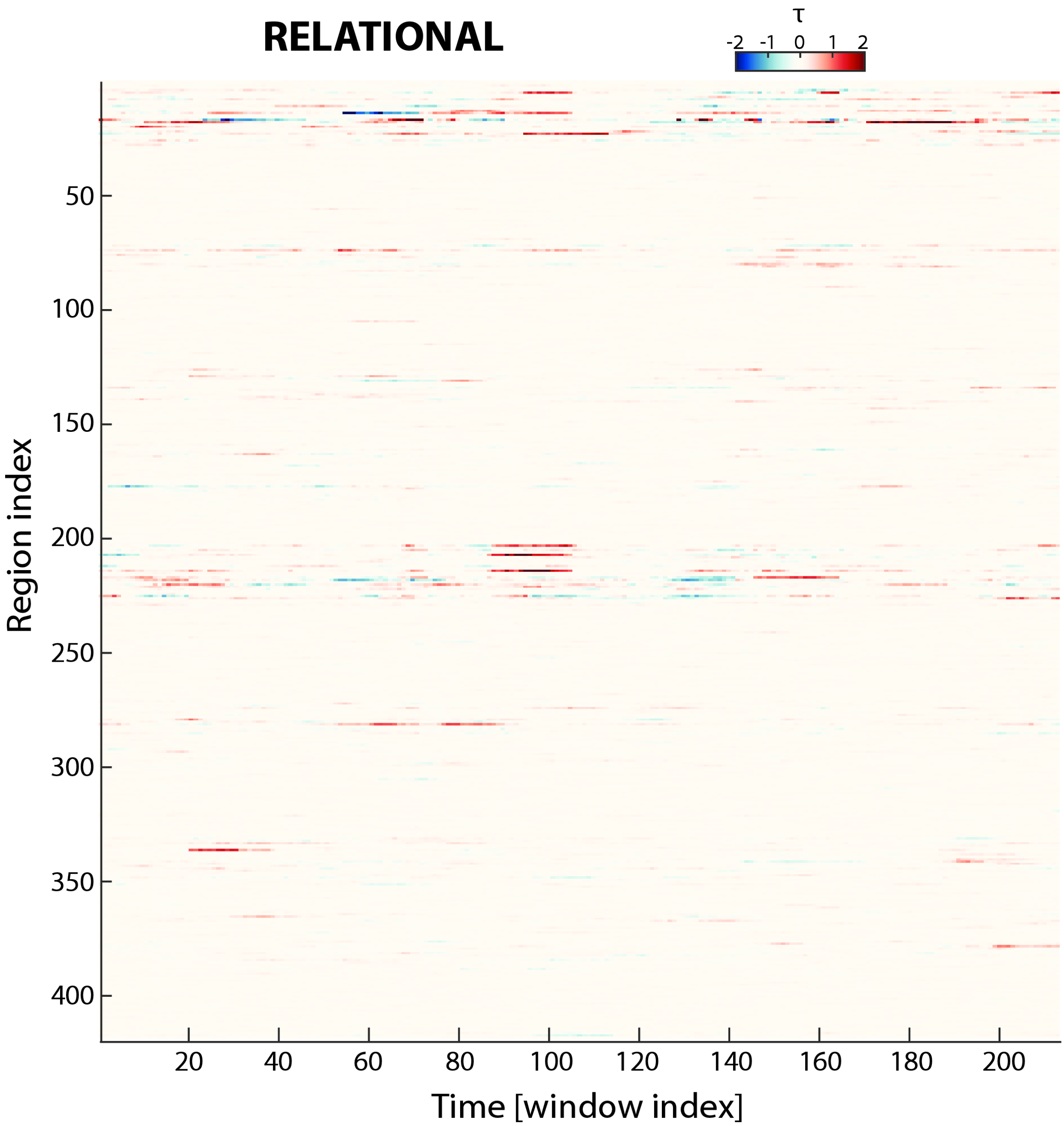


Figure 7: Evolution of causal effects during the relational task.

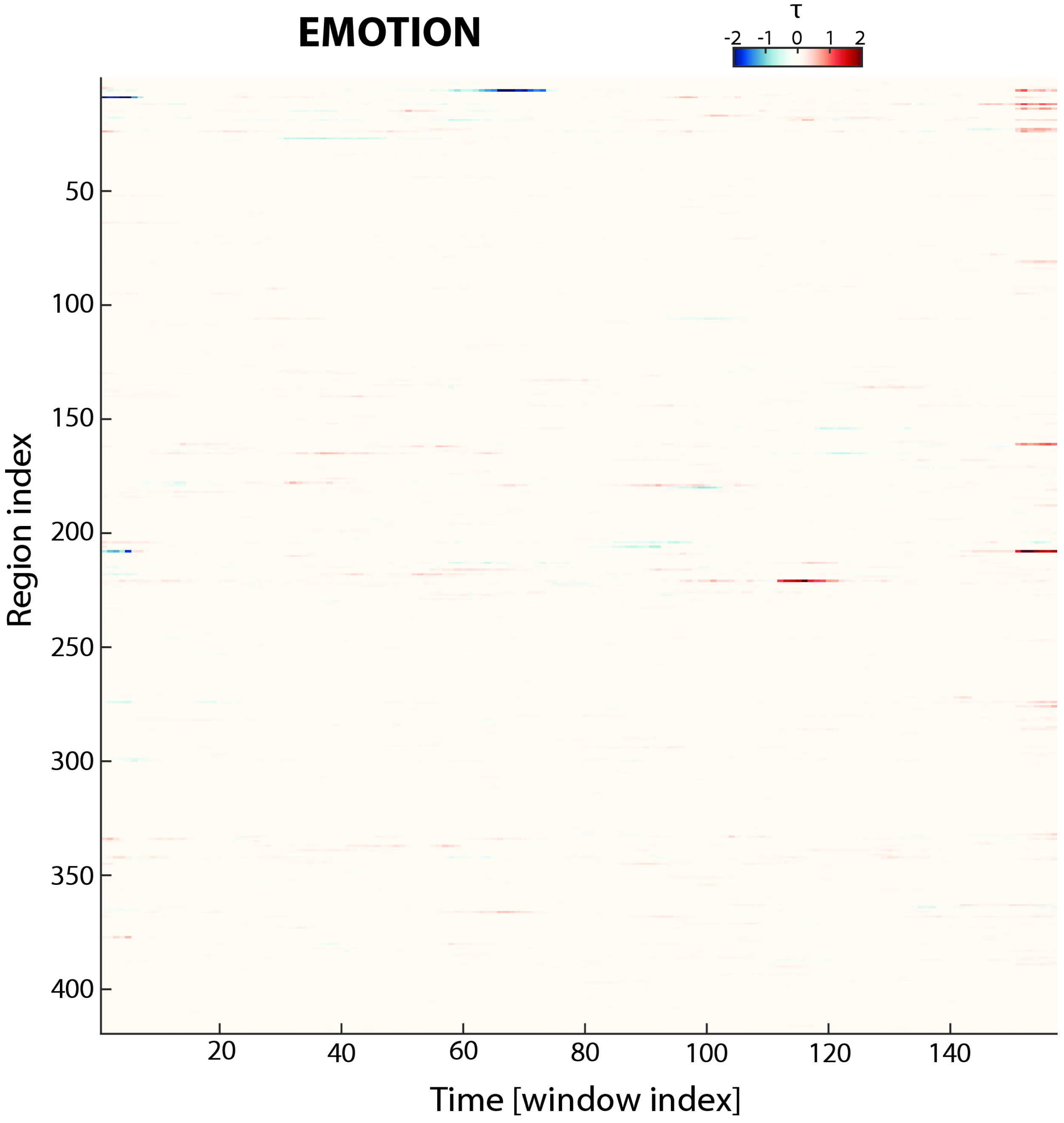


Figure 8: Evolution of causal effects during the emotion task.

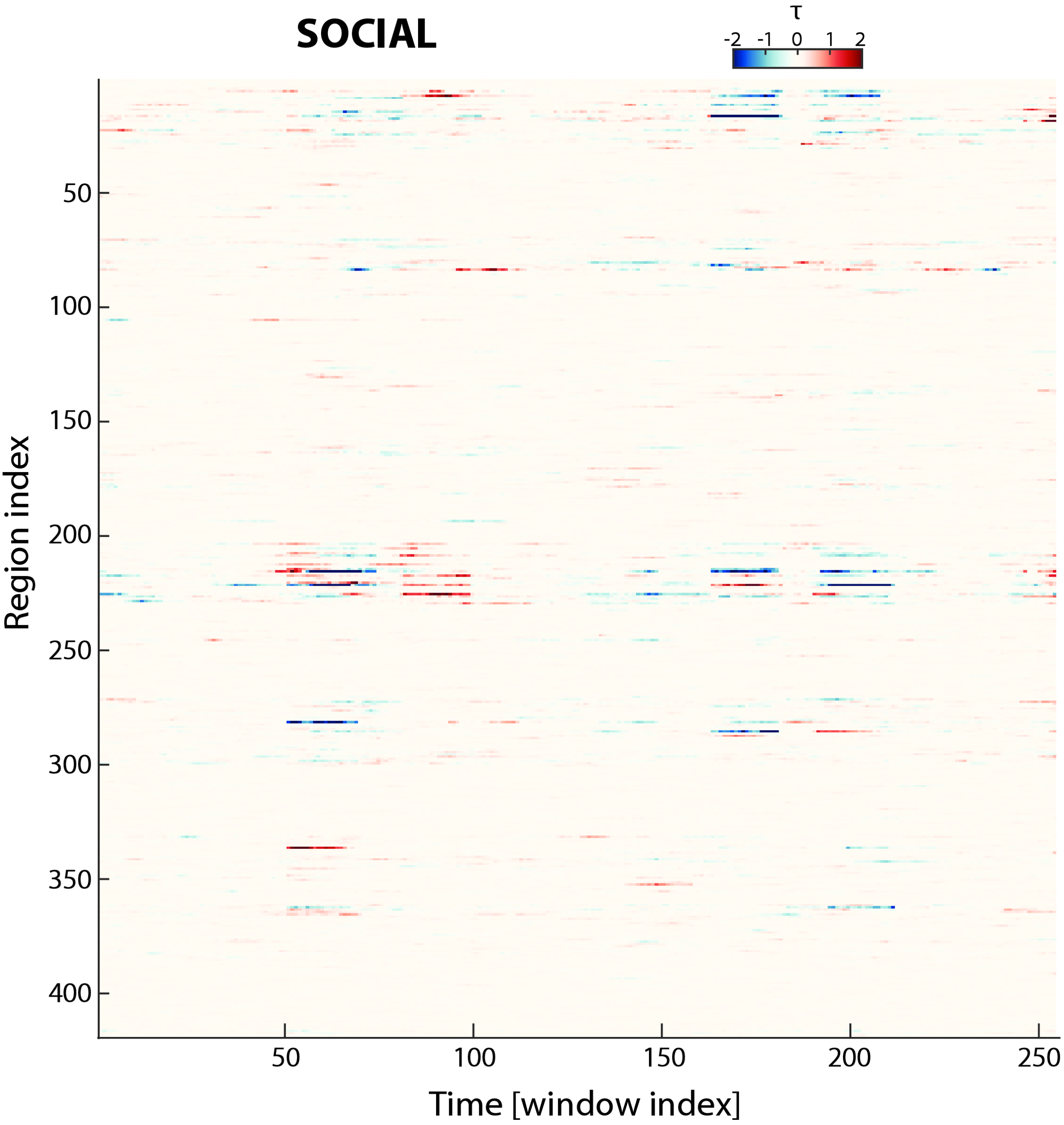


Figure 9: Evolution of causal effects during the social task.

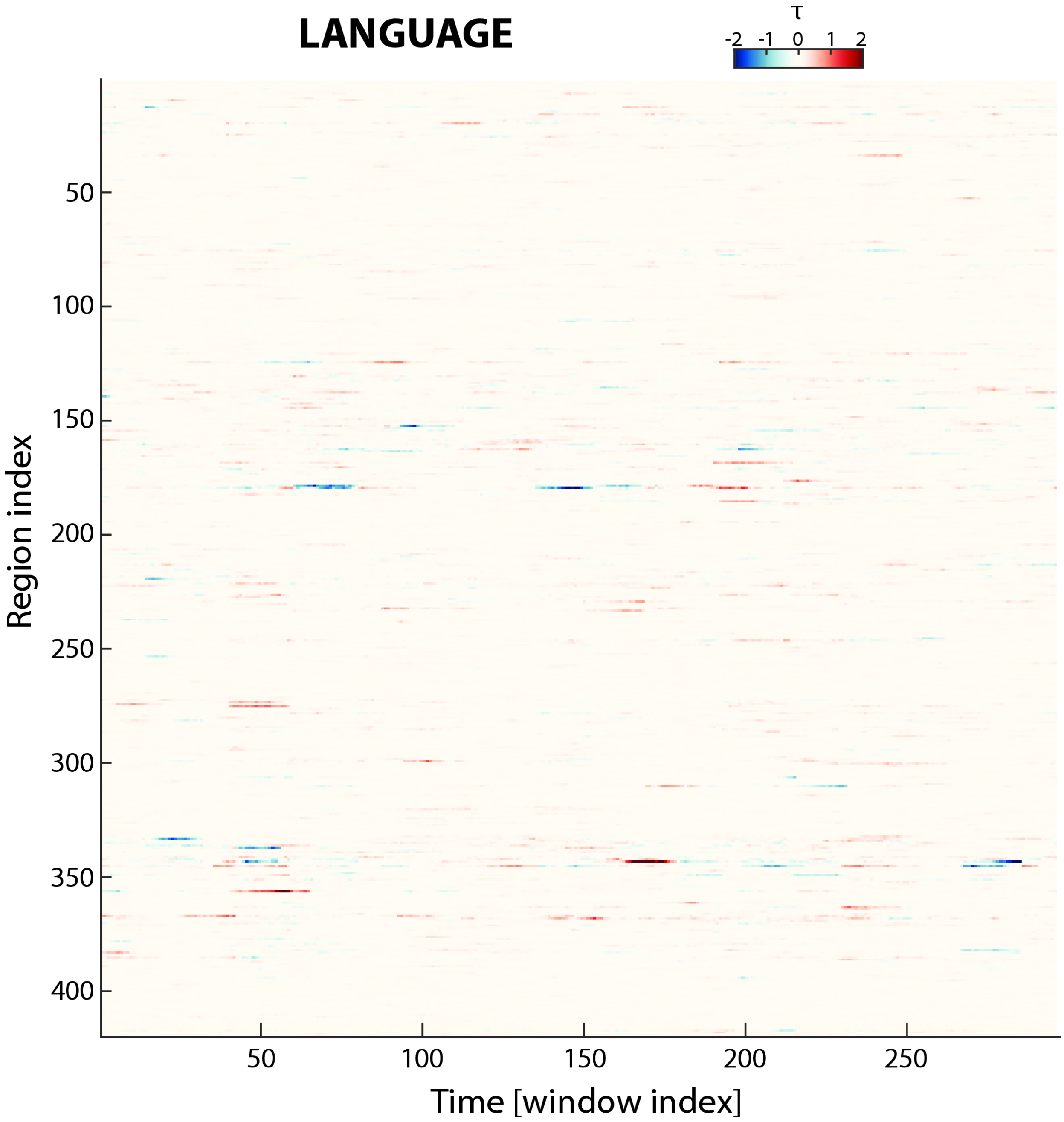


Figure 10: Evolution of causal effects during the language task.

*Impacts of run and (pre)processing choices on AoT patterns*

We performed additional analyses, for each of the investigated paradigms, in order to gauge the robustness of our findings to adjustments in our (pre)processing pipeline, or to the use of other input data. The assessed alternatives were the following:

1. The use of the right-left phase encoding direction recording as input data (*Run* variable)
2. The absence of global signal regression in preprocessing (*GSR* variable)
3. Instead of no censoring (case I), scrubbing of the preprocessed time courses (at a threshold of 0.5 mm framewise displacement[12]), removing only flagged samples (case II), also one sample before and two after each excised time point (case III), or three samples before and six after (case IV)
4. A different sampling scheme for the data points that enter $\tau$ computations, where instead of retaining all data points for a given subject (case I), $\frac{n_s^*}{S}$ samples were selected per subject taking the first available ones (case II), randomly picking them within the full recording (case III), or extracting a continuous block (notwithstanding excised volumes, if applicable) from a random starting location (case IV)
5. Another AoT measure, where non-normality is quantified using the Kullback-Leibler divergence between the error distribution of interest and a standard normal one (*Measure* variable)
6. For time-locked task paradigms, we also examined the differences between the use of full recordings, or of only task epochs (*Epochs* variable).

Stability of the results was quantified by Pearson's correlation coefficient between the AoT regional patterns obtained in each setting. To assess the impact of a given variable, we quantified

| | RS | MOTOR | WM | EMOTION | SOCIAL | LANGUAGE |
|---|---|---|---|---|---|---|
| *Epochs* | n.a. | 0.67 ± 0.07 | 0.38 ± 0.31 | 0.62 ± 0.11 | 0.41 ± 0.18 | 0.8 ± 0.09 |
| *GSR* | 0.37 ± 0.09 | 0.36 ± 0.09 | 0.45 ± 0.39 | 0.24 ± 0.13 | 0.58 ± 0.29 | 0.3 ± 0.07 |
| *Measure* | 0.75 ± 0.03 | 0.81 ± 0.04 | 0.75 ± 0.08 | 0.7 ± 0.07 | 0.81 ± 0.08 | 0.77 ± 0.05 |
| *Run* | 0.19 ± 0.07 | 0.19 ± 0.11 | 0.42 ± 0.29 | 0.15 ± 0.11 | 0.5 ± 0.28 | 0.19 ± 0.06 |

Table 1: Similarity between cases including the removal of baseline epochs or not, including global signal regression or not, considering a kurtosis-based or a Kullback-Leibler divergence-based AoT-sensitive metric, and asessing the left-right or right-left phase encoding run. Results are presented as mean ± standard deviation. RS: resting state. WM: working memory.

| | RS | MOTOR | WM | EMOTION | SOCIAL | LANGUAGE |
|---|---|---|---|---|---|---|
| I vs II | 0.74 ± 0.14 | 0.89 ± 0.09 | 0.87 ± 0.13 | 0.87 ± 0.14 | 0.92 ± 0.09 | 0.83 ± 0.12 |
| I vs III | 0.74 ± 0.18 | 0.94 ± 0.06 | 0.9 ± 0.12 | 0.93 ± 0.08 | 0.95 ± 0.07 | 0.88 ± 0.1 |
| I vs IV | 0.75 ± 0.18 | 0.94 ± 0.06 | 0.91 ± 0.11 | 0.93 ± 0.08 | 0.95 ± 0.08 | 0.88 ± 0.11 |
| II vs III | 0.73 ± 0.15 | 0.89 ± 0.09 | 0.88 ± 0.12 | 0.87 ± 0.14 | 0.92 ± 0.09 | 0.84 ± 0.12 |
| II vs IV | 0.73 ± 0.15 | 0.89 ± 0.09 | 0.88 ± 0.11 | 0.87 ± 0.14 | 0.92 ± 0.09 | 0.83 ± 0.12 |
| III vs IV | 0.75 ± 0.18 | 0.94 ± 0.06 | 0.91 ± 0.11 | 0.93 ± 0.08 | 0.94 ± 0.08 | 0.89 ± 0.1 |

Table 2: Similarity between different motion censoring schemes: no scrubbing (I), mild scrubbing (II), moderate scrubbing (III) and aggressive scrubbing (IV). Results are presented as mean ± standard deviation. RS: resting state. WM: working memory.

similarity when only this particular factor was varied, while all others were kept fixed. This yielded 64 values per variable, which we report below.

As can be seen from Table 1, regardless of the paradigm, both our original kurtosis-based measure and our alternative revolving around the Kullback-Leibler divergence yielded highly similar AoT patterns. The removal of baseline epochs had the largest influence on the social and working memory tasks. Whether to include global signal regression or not had a consistently sizeable effect in all paradigms, and so did selecting the first or the second available run.

From Table 2, it can be seen that AoT estimates remain extremely similar regardless of the extent of scrubbing applied to the data. This is strong evidence that head motion does not impact our results.

In Table 3, most sampling schemes can be seen to yield highly similar results, to the exception of case III (selection of the first time points for a given subject). This may be due to magnetization effects, or to physiological variables that would require a certain time to reach a steady state and

| | RS | MOTOR | WM | EMOTION | SOCIAL | LANGUAGE |
|---|---|---|---|---|---|---|
| I vs II | 0.59 ± 0.1 | 0.9 ± 0.07 | 0.85 ± 0.12 | 0.9 ± 0.08 | 0.9 ± 0.11 | 0.81 ± 0.09 |
| I vs III | 0.26 ± 0.11 | 0.29 ± 0.17 | 0.29 ± 0.3 | 0.32 ± 0.18 | 0.5 ± 0.22 | 0.29 ± 0.13 |
| I vs IV | 0.59 ± 0.11 | 0.81 ± 0.08 | 0.82 ± 0.15 | 0.75 ± 0.1 | 0.86 ± 0.12 | 0.75 ± 0.08 |
| II vs III | 0.26 ± 0.13 | 0.27 ± 0.18 | 0.3 ± 0.32 | 0.32 ± 0.18 | 0.51 ± 0.21 | 0.28 ± 0.14 |
| II vs IV | 0.61 ± 0.1 | 0.81 ± 0.08 | 0.83 ± 0.14 | 0.75 ± 0.1 | 0.85 ± 0.13 | 0.74 ± 0.08 |
| III vs IV | 0.24 ± 0.13 | 0.16 ± 0.19 | 0.3 ± 0.32 | 0.31 ± 0.19 | 0.44 ± 0.24 | 0.18 ± 0.13 |

Table 3: Similarity between different sampling schemes: all data for a subject (I), first samples per subject only (II), randomly selected samples per subject (III) and continuous block with random start per subject (IV). Results are presented as mean ± standard deviation. RS: resting state. WM: working memory.

would initially perturb the fMRI signals (*e.g.*, stronger heart rate fluctuations until one becomes at ease in the scanner).

In addition to the above, we also verified that the results were not affected by the use of a coarser ($R_2 = 219$ regions) or finer-grained ($R_3 = 819$ regions) atlas. As can be seen from Fig. 11, the extracted AoT patterns in the resting state and motor task cases remained similar regardless of atlas granularity.

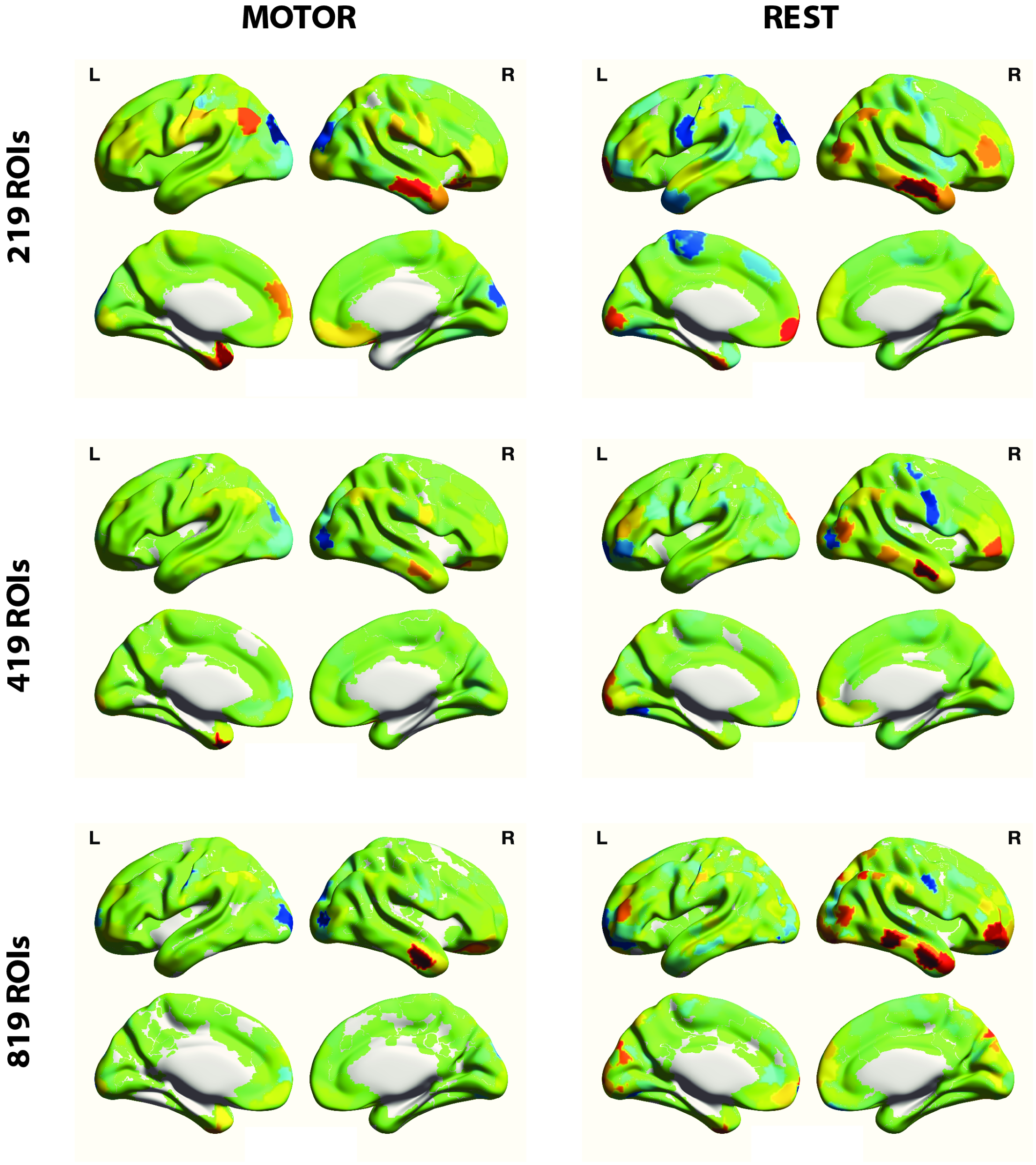


Figure 11: Regional AoT patterns obtained for the resting state and motor paradigms, using $n_s^* = 8000$ samples, when resorting to an atlas with 219, 419 or 819 regions of interest. ROI: region of interest.

### References


[1] Barch, D.M., Burgess, G.C., Harms, M.P., Petersen, S.E., Schlaggar, B.L., Corbetta, M., et al. Function in the human connectome: task-fmri and individual differences in behavior. Neuroimage 2013;80:169–89. doi:10.1016/j.neuroimage.2013.05.033.

[2] Kim, C., Kroger, J.K., Calhoun, V.D., Clark, V.P.. The role of the frontopolar cortex in manipulation of integrated information in working memory. Neuroscience letters 2015;595:25–29.

[3] Alnæs, D., Sneve, M.H., Richard, G., Skåtun, K.C., Kaufmann, T., Nordvik, J.E., et al. Functional connectivity indicates differential roles for the intraparietal sulcus and the superior parietal lobule in multiple object tracking. Neuroimage 2015;123:129–137.

[4] Xu, Y.. The role of the superior intraparietal sulcus in supporting visual short-term memory for multifeature objects. Journal of Neuroscience 2007;27(43):11676–11686.

[5] Christoff, K., Prabhakaran, V., Dorfman, J., Zhao, Z., Kroger, J.K., Holyoak, K.J., et al. Rostrolateral prefrontal cortex involvement in relational integration during reasoning. Neuroimage 2001;14(5):1136–1149.

[6] Farrer, C., Frey, S.H., Van Horn, J.D., Tunik, E., Turk, D., Inati, S., et al. The angular gyrus computes action awareness representations. Cerebral cortex 2008;18(2):254–261.

[7] Guidali, G., Pisoni, A., Bolognini, N., Papagno, C.. Keeping order in the brain: the supramarginal gyrus and serial order in short-term memory. Cortex 2019;119:89–99.

[8] Martinez-Trujillo, J.C., Cheyne, D., Gaetz, W., Simine, E., Tsotsos, J.K.. Activation of area mt/v5 and the right inferior parietal cortex during the discrimination of transient direction changes in translational motion. Cerebral Cortex 2007;17(7):1733–1739.

[9] Hartwright, C.E., Apperly, I.A., Hansen, P.C.. Representation, control, or reasoning? distinct functions for theory of mind within the medial prefrontal cortex. Journal of Cognitive Neuroscience 2014;26(4):683–698.

[10] Fletcher, P.C., Happe, F., Frith, U., Baker, S.C., Dolan, R.J., Frackowiak, R.S., et al. Other minds in the brain: a functional imaging study of "theory of mind" in story comprehension. Cognition 1995;57(2):109–128.

[11] Theiler, J., Eubank, S., Longtin, A., Galdrikian, B., Farmer, J.D.. Testing for nonlinearity in time series: the method of surrogate data. Physica D: Nonlinear Phenomena 1992;58(1):77–94.

[12] Power, J.D., Barnes, K.A., Snyder, A.Z., Schlaggar, B.L., Petersen, S.E.. Spurious but systematic correlations in functional connectivity mri networks arise from subject motion. Neuroimage 2012;59(3):2142–2154.
